# Pairwise interactions, feedback rule changes, and deliberative decisions underlie honeybee inflight group coordination

**DOI:** 10.1101/2024.10.10.616169

**Authors:** Md. Saiful Islam, Imraan Faruque

**Affiliations:** School of Mechanical and Aerospace Engineering, Oklahoma State University, Stillwater, Oklahoma, USA

## Abstract

Systematic descriptions of the underlying interaction rules that insects use to support group and swarm flight has the potential to contribute to mathematics, biology, and robotics, including aerial swarming under sensory and computational limitations. This study analyzes 1,000 trajectories of flying honeybees in crowded conditions approaching a moving stimulus and finds how during this stimulus, honeybees coordinate flight through pairwise interactions involving a novel three-zone decision-making process. The experimental setup consists of 3-D position reconstructions via a high speed camera system recording honeybee foragers returning to a hive entrance actuated to move robotically. The analysis consists of neighborhood identification through three methods (cross-correlation, distance threshold, and average distance threshold), which reveals the dominant interaction is pairwise. The individual leader-follower pair interactions are then tested against three regulation candidates: optic flow, relative velocity, and optical expansion rate, based on minimizing root mean square error. The results show that each follower demonstrates a three stage process involving a feedback rule change, linked by an intermediate observation/decision phase. During the initial “lock” phase, an insect maintains a consistent optical expansion rate until inter-agent distance closes to 10 cm. The regulation candidates then undergo large variations during a relatively long observation/decision zone, with 1.04 seconds being the average time in the decision zone. 79% of the paired insect entries into the decision zone result in subsequent re-engagement to track the same initial leader, while 21% result in disengagement from the group behavior. Visual regulation candidate comparison in the third stage indicates that upon re-engagement, the follower relative velocity is regulated to provide consistent velocity matching between agents. The third stage’s velocity tracking is consistent with a closed-loop feedback proportional-integral (PI) controller regulating velocity tracking error. Across the insect population studied, the proportional gain remained showed minimal variability over individuals, a derivative gain was considered and found negligible, and the integral gain varied by individual. Collectively, these findings underscore the existence of an alternative swarm architecture, highlighting individual decision-making capabilities, feedback regulation target changes, and the presence of reactive, deliberative, and moderate (PI control) timescale interaction rules contained within aerial groups.

## Notation

***W*** Adjacency matrix

*V* Set of nodes of the graph

*ϵ* Set of edges of the graph

***L*** Laplacian matrix

***M*** Degree matrix

***N*** Number of insects

*τ* Lag time of cross correlation

*T* Time length of trajectory

*S*_*i,j*_(*τ*) Cross-correlation between insects *i* and *j*

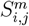 Maximum cross-correlation value from *S*_*i,j*_(*τ*)

*G* = {*ĝ*_*x*_, *ĝ*_*y*_, *ĝ*_*z*_} Fixed (inertial) frame

***X***_*i*_(*t*) = [*x*_*i*_(*t*), *y*_*i*_(*t*), *z*_*i*_(*t*)]^*T*^ Agent *i* position vector in *G* frame

*y*_*i*_(*t*) *y*-coordinate of the insect *i* in *G* frame

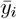 Temporal average *y*-coordinate of the insect *i* in *G* frame

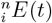 Euclidean distance between agent *i* and agent *n*

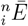 Temporal average Euclidean distance between insect *i* and *n*

*γ* Distance criteria for group determination

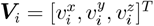 Velocity vector of insect *i* in *G* frame

***v***_*i*_ Unit vector of the velocity vector of insect *i*

*R*_*o*_(*t*) Optical expansion rate candidate

*R*_*f*_ (*t*) Optic flow candidate

*R*_*v*_(*t*) Relative velocity candidate

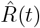 Ideal magnitude of any visual candidate

*R*_*T*_ (*t*) Set of three visual regulation candidates

*p*(*t*) Slope of a line

RMSE Root mean square square error

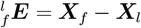 Relative position vector between a follower and leader in *G* frame

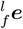 Unit vector of relative position between a follower and leader

*θ* Heading heading angle between follower *f* and leader *l* points

*σ*(*t*) Unsigned angle of individual motion

*Ψ*(*t*) Clockwise or anticlockwise sign of individual direction of motion

*σ*_*fl*_(*t*) Unsigned angle between follower and unit vector of relative position

*Ψ*_*fl*_(*t*) Sign of relative direction of motion

Δ*β*_*fl*_(*t*) Signed relative direction of motion

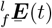 Forced mapped coordinates’ vector of paired follower and leader

Δ*t* Sampling time step

*F* Length of interacting domain for the force map

(*q, r*) Length of each row and column bin of the force map

*k*_*p*_, *k*_*i*_, *k*_*d*_ Proportional, integral and derivative gain

***E***_*r*_(*t*) Error between true and simulated velocity

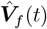 Simulated follower velocity

FIT Fit percentage of simulated response PDF Probability density function

*J*(*x*, 0) PDF of *x* coordinate of followers in force map

*J*(0, *y*) PDF of *y* coordinate of followers in force map

*J*(*x, y*) Joint PDF of *x, y* coordinates of followers in force map

### Sub/super scripts

_*i*_ agent index, *i* ∈ [1, *N*]

_*f*_ follower agent

_*l*_ leader agent

## Introduction

‘Individual insects in flying insect groups maintain coordination with their neighbors despite tight constraints on neural resources, including on sensory structures and in-flight communication. Many existing engineering swarm constructions (such as the well-known Boids model) rely on agents interacting with three effects: long range attraction, mid-range velocity alignment, and short range repulsion or collision avoidance. These agent interaction rules have been shown theoretically to support coordinated behaviors similar to natural systems like fish schools, bird flocks, and insect swarms (Reynolds, 1987; Cucker and Smale, 2007). Control theoretic constructions often leverage an engineered separation of guidance, navigation, and control (GNC) disciplines, while robotic approaches involve sensing, perception, decision, and action (SPDA) paradigms (such as “sense, decide, engage” or “observe, orient, decide, act” (OODA) in military contexts). However, agents seen in biological groups incorporate more individually diverse constructions involving more varied roles, reaction strengths, and individual goals. Rather than adhering to GNC, SPDA, OODA frameworks implementing attraction/repulsion or cohesion/alignment/avoidance models, local individual models may incorporate more diverse behaviors such as homing, neighbor tracking, observation, or foraging. Individual coordination rules can also vary over organismal development, such as velocity alignment appearing before attraction in schooling fish (Paz et al., 2023). Homogeneous swarm models relying on current engineered paradigms do not fully describe these biological collectives. These challenges have motivated research into how individual agent roles and differences can affect group behaviors. An individual agent’s connectivity, particularly its choice of which neighbors are included in a particular agent’s interaction neighborhood, has ramifications on group performance, individual sensing and feedback, and an implementation’s computational demands. The need for systematic tools to extract the underlying interaction network from measured data has complicated in-flight local interaction network analysis. Correlation, transfer entropy, causal entropy, and Granger causality have been applied to this reconstruction problem, which are systematic ways to provide successive pairwise analysis. The differing interaction network sizes across species may reflect the availability of sensing, feedback, and neural resources as well as the ecological context of the behavior.

This study measured the three-dimensional trajectories of honeybees in dense flight as they approached their hive entrance as the entrance location is actuated dynamically. It first considers what neighborhood size insects maintain awareness, in terms of the number of agents each individual insect in the group receives information from (e.g., its in-degree in an information flow network). This study reconstructed the implicit information flow interaction network graph using correlation and distance-based methods. The reconstructed information graph showed insects responded to a maximum of 2-4 neighbors, with 28% of all recorded group members being included in a pairwise interaction. The relative dominance of pairwise interactions over other interaction sizes motivated a focus on these pairwise interactions, or “leader-follower” pairs. The analysis of these leader-follower trajectories identified optical expansion rate as the parameter that the follower initially regulates during its approach.

This study establishes for the first time a three-zone structure of individual behaviors across this pairwise connection. The zones consist of (a) regulating the optical expansion rate, (b) a lengthy observation period supporting deliberative decision-making regarding the third zone, and (c) either disengagement or velocity matching relative to the leader using a proportional-integral feedback rule. In the decision-making zone, the insect did not regulate relative to the moving visual targets in its observation field. However the majority of trials showed a re-engagement, and the same leader was preserved. The velocity-matching behavior fit a proportional integral derivative (PID) controller structure with a constant proportional gain and a integral gain that varied over individuals.

The overall study flowchart is illustrated in Fig. 1, showing the biological flight experiment as the first block and the neighborhood analysis as the second block, which includes three independent analysis approaches (cross-correlation, inter-agent distance threshold, and pairwise insect analysis). The leader-follower pairs were then isolated, and the final block shows the study component that developed the three zone model. This study includes a hypothesis-based (H1, H2, and H3) isolation of individual feedback rules, and the transition criteria between zone 1 and 2.

**Figure 1:**
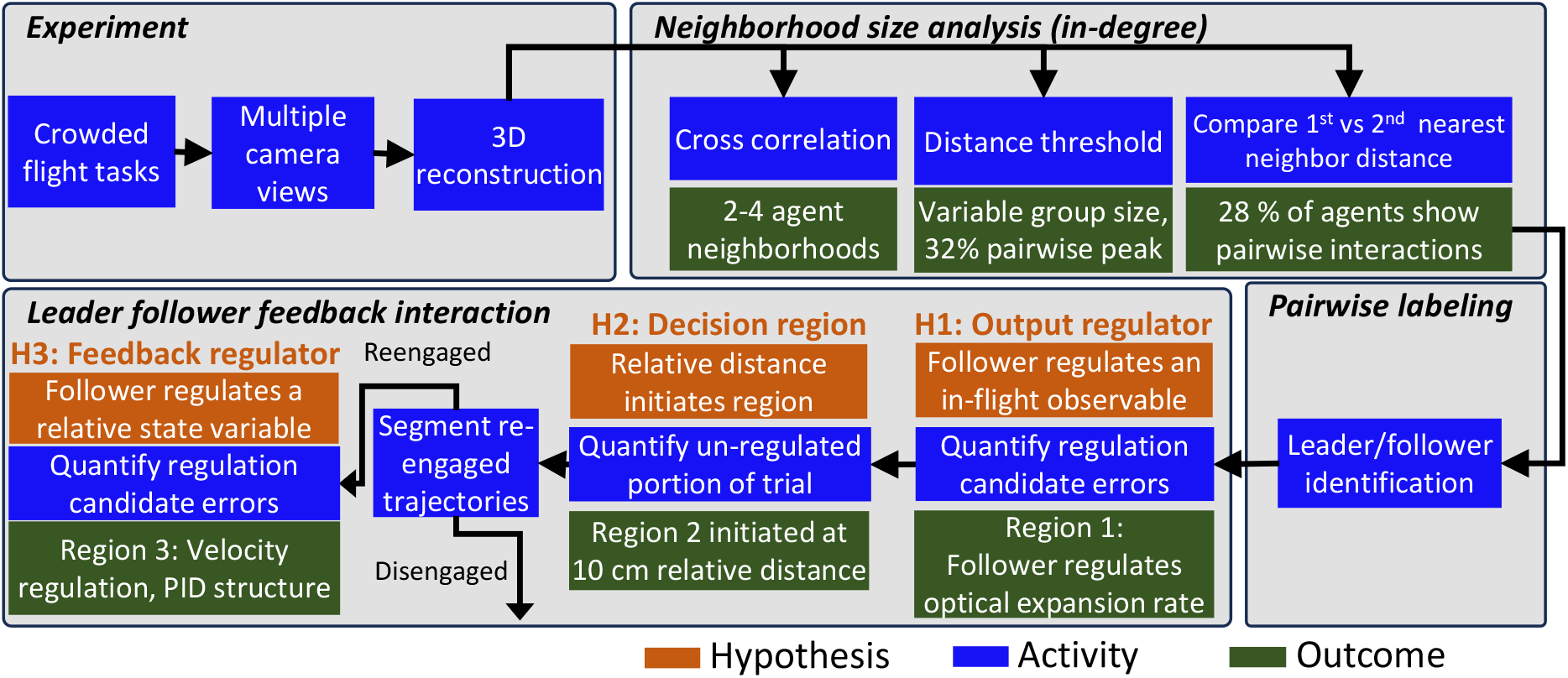
Flowchart outlining the overall study workflow, with H1, H2, and H3 denoting hypotheses, blue blocks describing activity, and green blocks representing the results.

### Previous works review

#### Coordination rules

Discovering the coordination rules than underlie natural collective behaviors has received significant scientific attention. Natural swarms are of particular interest as an example of a cohesive self-organized non-equilibrium collective behavior that lacks a clear order parameter (Vicsek and Zafeiris, 2012; Lopez et al., 2012; Attanasi et al., 2014). Previous work examined swarm cohesion from the perspective of metric distance (AOKI, 1982; Huth and Wissel, 1992; Couzin et al., 2002), network topology (eg, position rank (Ginelli and Chaté, 2010)), apparent visual size (Lemasson et al., 2009, 2013), and agent-relative spatial distribution Gautrais et al. (2012). These modeling approaches may work to create a bio-inspired description or incorporate more contemporary engineering constructions (Bode et al., 2010, 2011). Entropy based analysis (Lord et al., 2016), information theoretic transfer (Schreiber, 2000), and time series correlation (Puckett et al., 2015) methods have made progress in describing this behavior. However, correlation analysis’ directionality and linearity restrictions, sample size impacts, and need for outlier consideration limits its ability to describe this behavior and complicates interpretation (Papana, 2021). Similarly, information and entropy-based analysis does not identify the path of information flow; considering a time delay can help but does not explicitly identify information exchanged from a shared input signal (Shannon, 1948). However, much remains to be discovered about the mechanism and function underlying biological systems’ coordination in groups, particularly when supporting high bandwidth physics tasks with limited neural systems and connectivity.

#### Agent neighborhood size

The number of neighbors that an individual agent attends to is a practical question which has been first addressed by using correlation analysis to provide an upper bound on the interaction neighborhood (Sun et al., 2014; Sun and Bollt, 2014). Scale-free correlations in birds indicate that spatial correlation is not constant, and instead scales with the linear size of the flock (Cavagna et al., 2010) and the actual neighbor interaction range is not adequately described by correlation analysis (Cavagna et al., 2010). The number of interaction neighbors has also been estimated via anisotropy analysis. Anisotropy is computed via the average orientation change of a given agent’s neighbor, an analysis that indicates flocking starlings maintain 6-7 pairwise connections with a fixed number of birds (Ballerini et al., 2008) (a topological limit), weakening support for a fixed metric distance-based hypothesis such as the sensing or interaction radius used in many engineered distributed systems (Czirók and Vicsek, 2000). Small amounts of measurement noise can destroy cohesion in Vicsek self-propelled flocks, an effect which is reduced by including long-range alignment Zumaya et al. (2018) even when only a few long-range interactions are present. Interaction with one to two neighbors at a time is also seen in the short-term directional correlation of U-turn events *Hemigrammus rhodostomus* fish groups (Jiang et al., 2017). The limited neural resources and comparatively poor optics (relative to these vertebrate examples) suggest the possibility that an individual insect’s interaction neighborhood may be more tightly constrained, but a parallel measurement of the neighborhood size in insect swarms has not yet been experimentally conducted.

#### The role of pairwise connections in collective motion

Coordinated behavior can arise from low agent network sizes, including when the agents maintain only a single interaction neighbor–or a pairwise connection. Previous studies have found pairwise interactions in flying birds, bats, and pigeons, showing the benefits of these coupled flights. Marching locusts’ coordinated behaviors emerge as a result of their paired relationships that facilitate information transfer without the need for outside perturbation (Buhl et al., 2006). *Anarete pritchardi* swarm’s collective motion has not been clearly described beyond frequent pairwise harmonic oscillations embedded in a less structured environment (Okubo and Chiang, 1974). Midge swarm movements show two distinct modes: a low-frequency (independent) mode and a high-frequency mode associated with pairwise interaction (Puckett et al., 2015). Insect pair-formation includes long-range sound-based attraction forces, with attraction increasing at the center of the swarm when sound is louder and decreasing at its boundary where sound is weaker (Gorbonos et al., 2020). Two biological goals might support this pairwise formation: to explore the volume of swarms for potential predators or search for female partners (Downes, 1969; Neems et al., 1992; Puckett et al., 2015). Leader-follower behaviors have also been reconstructed using time delay correlation in bat and pigeon flights. Bird species that form persistent pair bonds may retain the pair bond during inflight coordination, reducing individual attentional workloads (Ballerini et al., 2008; Billah and Faruque, 2024), and persistent paired interactions in flocks can increase the flock’s separation-resistance while navigating obstacle fields and lower the attentional requirements of the non-paired agents (Jolles et al., 2013; Ling et al., 2019a; Nagy et al., 2019). Pairwise bird flight analysis compared paired and non-paired birds to find that unpaired birds show quicker changes in their inter-agent distance than paired birds, wing beat frequency is lower for paired birds, and the anisotropic delay is lower in paired birds (Ling et al., 2019a). The leader-follower network was reconstructed by a pairwise time-delayed angle correlation from pigeon flocks and bat flocks (Nagy et al., 2013; Giuggioli et al., 2015). Fish that mimic their neighbor’s headings may modulate how long they imitate others, which helps in the formation of a stable flock and increases their locomotion capacity (Agrillo et al., 2008). Despite this progress, the interaction rules that enable flying animals to demonstrate efficient pairwise measurements in flight remain an open research question.

#### Insect feedback control laws for inflight pairwise/interception

Research on insects, fish, bats, and humans demonstrated the widespread use of constant bearing strategies during target pursuit. In this strategy, an animal maintains a constant angle between its heading and the target (Olberg et al., 2000; Lanchester and Mark, 1976; McBeath et al., 1995; Fajen and Warren, 2004; Wardill et al., 2017; Ghose et al., 2006). In teleost fish, previous research found *Acanthaluteres spilomelanurus* fix their eyes on a target while their body continues forward motion, generating predictive pathways while pursuing a moving target. A linked control system has been proposed to explain the role of eye and body movements in generating these predictive pathways Lanchester and Mark (1976). Constant absolute target direction is an alternate strategy in which an animal adjusts its flight direction, which has been observed in echolocating bats in capturing flying prey. This process, typically completed in less than one second for detection and tracking, was analyzed through infrared high-speed videography and delay differential equations to understand the complex pursuit trajectories. The constant absolute target direction strategy has drawn analogies to proportional-navigation (PN) missile guidance (Ghose et al., 2006). Dragonfly interception strategies may also involve the use of three sensorimotor rules: initiating a head saccade towards the potential prey within approximately 50 milliseconds; assessing the angular size and angular speed of the prey; and executing an interception flight while visually fixated, lasting around 300 milliseconds (Lin and Leonardo, 2017). Robber flies (*Laphria*) generate an interception course by fixing their bearing angle while approaching a prey within 29 cm distance and invariant image properties or distance estimation have been suggested as the possible explanations of this constant bearing angle (Wardill et al., 2017). Visual reconstruction during mating flights indicates that honeybee drones continuously modulate their heading rate to align their bodies with the drone-queen axis using optical feedback (Gries and Koeniger, 1996). Honeybees may also respond optically to approaching threats such as wasps (Tan et al., 2012, 2013), and visual reconstruction during mating flights indicates that honeybee drones continuously modulate their heading rate to align their bodies with the drone-queen axis using optical feedback (Gries and Koeniger, 1996). However, these studies did not extract the in-flight feedback and decision architectures used between insects navigating aerial groups of their peers.

#### Decision-making capabilities of insects

Insects exhibit individual decision-making during forage target choice, measurable based on the speed and accuracy of solo insects landing on a forage target, indicates that flower selection is influenced by the flower color and smell, flower species, temperature, individual insect health or colony health (Gold and Shadlen, 2007; Khan et al., 2021; Chittka, 2022; MaBouDi et al., 2023). Honeybees also demonstrate collective decision-making in nest site selection, where accurate decision-making requires only a minority of agents. Comparisons with artificial voting models that draw analogies to human group decision-making have suggested the speed and decision time variation of individuals may play a critical role in a group decision (Bell, 2008; Ulyshen et al., 2023), including supporting a value-sensitive bifurcation (Gray et al., 2018). These analogies between social insect colonies and the human neural systems has inspired constructions in which insects (neurons for the brain) provide possible options, and a decision is made when combined individual agreements pass a threshold dependent on the accuracy and speed of the decision (Marshall et al., 2009). The effect of sample time on insect decision time indicates honeybees make accurate acceptance decisions more quickly than incorrect ones, and monkey decision-making experiments show similar results (Churchland et al., 2008). Conversely, human decision accuracy decreases with response time increases (Murphy et al., 2016), and during the decision-making periods, animals primarily gather information from the surroundings in a short time window (Thura et al., 2012). Robberflies incorporate predictive saccades and head fixation into their gaze control to support aerial predation. The fixation provides an average of 72.2 ms of observation and decision window (Talley et al., 2023).

Locust swarm marching shows pause and go motion, suggesting periodic assessment. Experimental and simulated analysis has suggested that animals frequently assess their surroundings by intermittent motion, a process that may be regarded as individual decisions about whether or not to join a swarm (Ariel et al., 2014). Unfortunately, due in part to a lack of studies involving in-flight group decision-making experiments, the influence of neighbors on individual insect decisions during flight is still not well understood.

### Contribution of this study

Despite the progress in understanding coordination rules, much remains unknown about how biological systems achieve their coordination, particularly during high bandwidth team tasks under limited neural and communication resources. Although agent neighborhoods and network connectivities have been quantified for larger biological systems, insects’ limited neural resources and comparatively poor optics (relative to these vertebrate examples) suggest the possibility that an individual insect’s interaction neighborhood may be more tightly constrained, but a parallel measurement of the neighborhood size in insect swarms has not yet been experimentally conducted. In particular, the pairwise interactions seen in flying bird groups have not been documented in honeybees, and the aerial pursuit strategies in this eusocial insect species are limited to mating or predation studies, not forager-to-forager team interactions. The inflight feedback laws used in these group member interactions remain unquantified. In particular, inflight decision-making has focused on discrete changes, e.g. instantaneous changes in target towards a higher value target, target abandonment, or feedback law change. The individual decision making and agent selection criteria architectures they implement are not yet clear, particularly the observation, processing, and deliberation timescales. The existence of a lengthy in-flight observation or deliberation period leading to a subsequent decision has not yet been established in flying insects.

The major contributions of this study include quantifying in-flight neighbor-to-neighbor information flow in honeybees flying in an outdoor experimental crowded task, extracting the implicit information flow paths, regulation candidates and feedback laws, and discovering an intermediate un-regulated decision period. More specifically, this study

1. measured experimental flight trajectories for multiple honeybees during crowded hive entrance approaches
2. identified the interaction network size for individual agents, including using parallel methods to passerine bird flocking studies. The analysis revealed small network sizes in insect groups in which the most common interaction is pairwise (i.e., a single neighbor)
3. determined the optical signal that an agent initially regulates to coordinate its flight with a neighbor agent, showing the regulated variable is the newly defined quantity “optical expansion rate”
4. discovered a 3-zone pair-wise interaction behavior including initial tracking, a disengaged un-regulated period, and a post-decision re-engagement period
5. quantified the time period spent in the unregulated decision period, showing that its length and re-engagement suggests a consistent observation that supports a deliberative decision versus the instantaneous change between two regulation rules previously
6. showed that the re-engaged regulation behavior is a velocity regulation consistent with a proportional-integral (PI) controller structure.

## Results

### Neighborhood analysis

Many animals, including birds and fish, incorporate information from only a subset of their neighborhood for their inter-agent communication, which may often be a single neighbor (i.e., a pairwise interaction), as reviewed in the previous works section. This study reports on an outdoor experiment to capture visual tracking trajectories to investigate the interaction neighborhood of honeybees, as shown in Fig. 2 and Methods: Insect stocks and handling. From individual insect trajectories captured by the imaging system (Method: Imaging system), the underlying interaction network was first reconstructed by applying graph theoretic correlation range analysis (Refer to Methods: Correlation range). An illustration of 2D and 3D trajectories of a group of six insects and their associated graph are shown in Figs. (3b-3c) and S1 Video. Over 1000 groups of flying insects, over 70% of insects (nodes) showed 2-4 connections (edges) to their neighbors, as seen in Fig. 3h. A second method, a distance-based threshold approach with varying thresholds was applied to segment insects in group flight into pairs, triples, quadruples, or larger-size groups.

**Figure 2:**
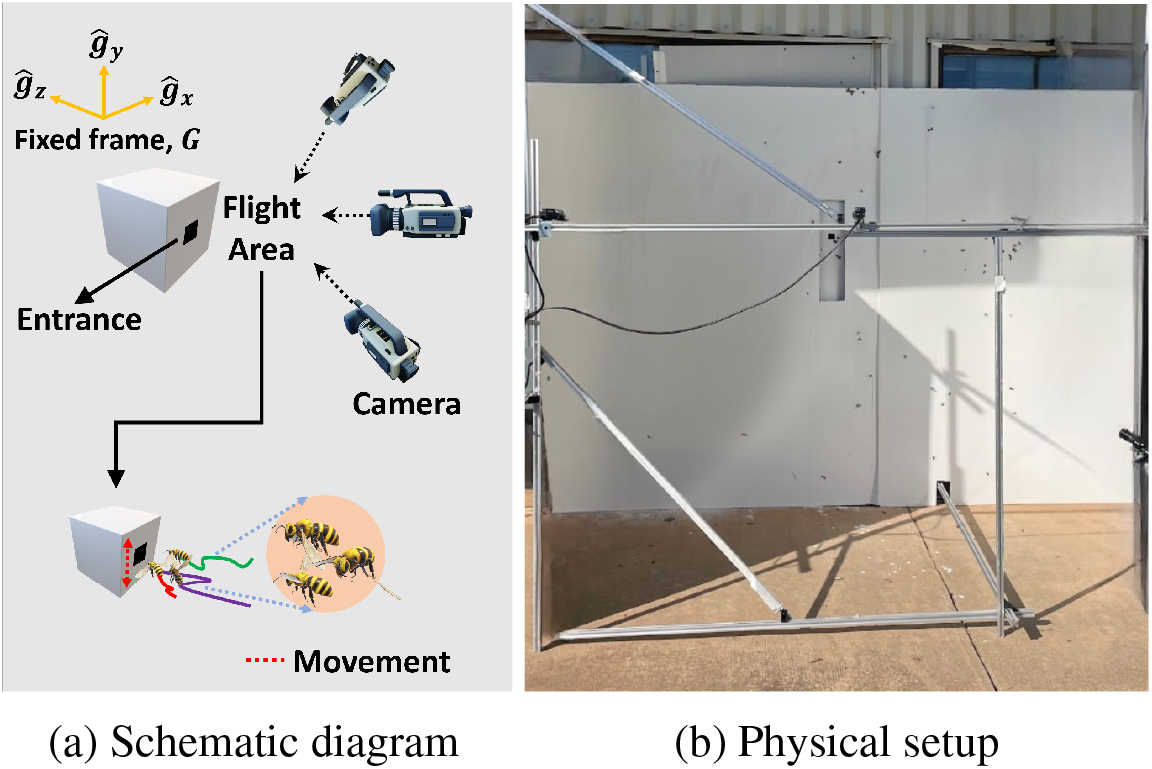
The experimental design includes a moving stimulus (a) surrounded by an outdoor multi-camera visual tracking system (b) which records the trajectories of insects as they approach the entrance in varying group sizes and stimulus conditions.

**Figure 3:**
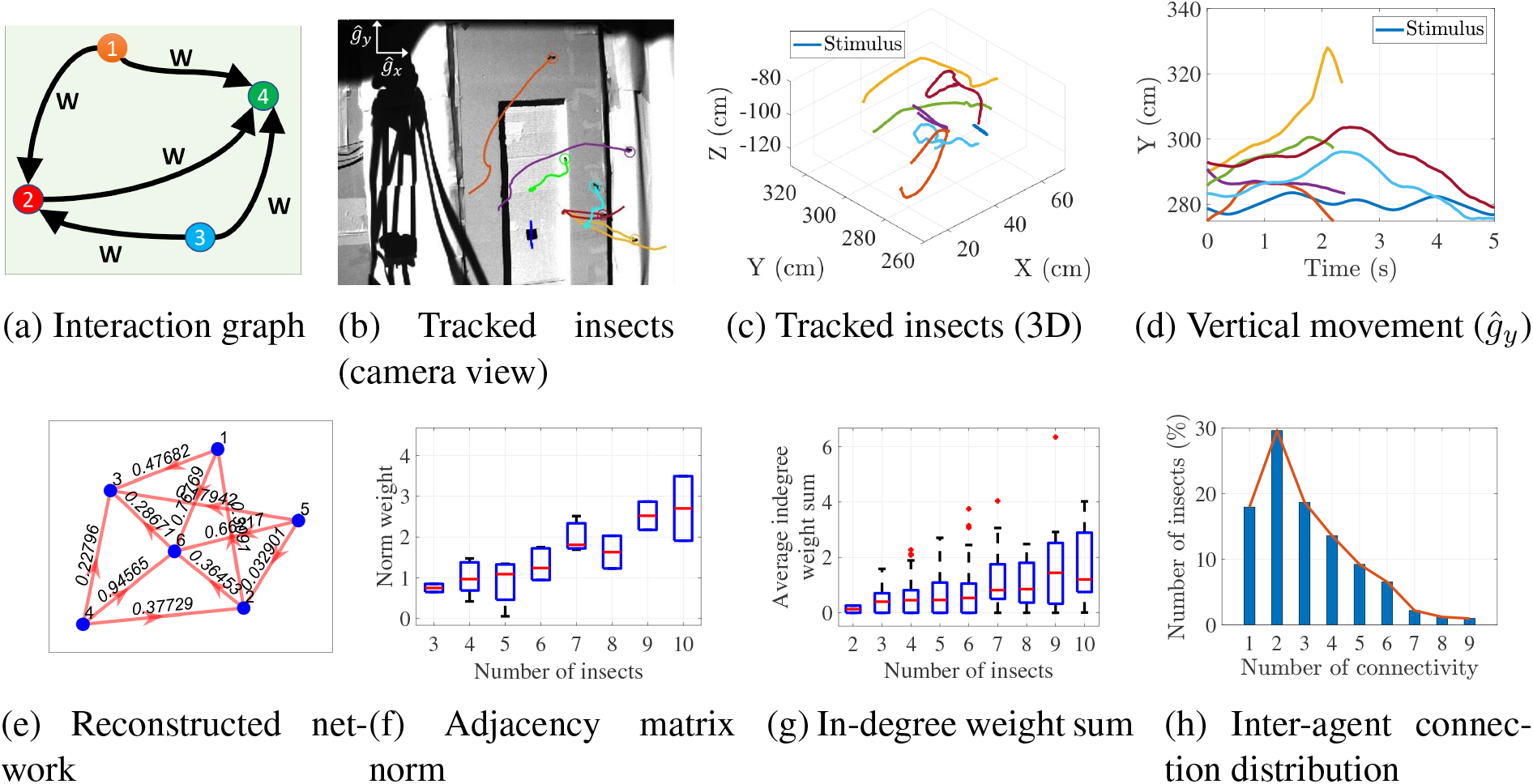
The implicit visual communication may be drawn as an information flow network having edge weights *w*_*ij*_. For an example of 6 insects approaching the hive (b), the insect motions were reconstructed in 3D (c) and the vertical movement (d) considered to identify the information flow. The reconstructed network (e) showed an increase in information flow with group size, and the in-degree (g) also grew. The connection distribution (h) shows that the most common interaction is pairwise, with 30% of insects being involved in a pairwise connection and 80% of agents being connected to 1-4 neighbors.

This analysis isolated formations by calculating the temporal average of the inter-agent distance of each insect *i* to its neighbors 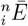, where *n* is the neighbor from the set [1…*N* − 1], and sorted them by a threshold *γ* (refer to Methods: Group sizes sorted by distance threshold). Figure. 4a shows the effect of the threshold variation on the group formation. The pair and triple formation percentages peak at *γ* = 7 cm and at *γ* = 10 cm, respectively, after which the relative fraction of quadruple and pentuples continues to increase.

**Figure 4:**
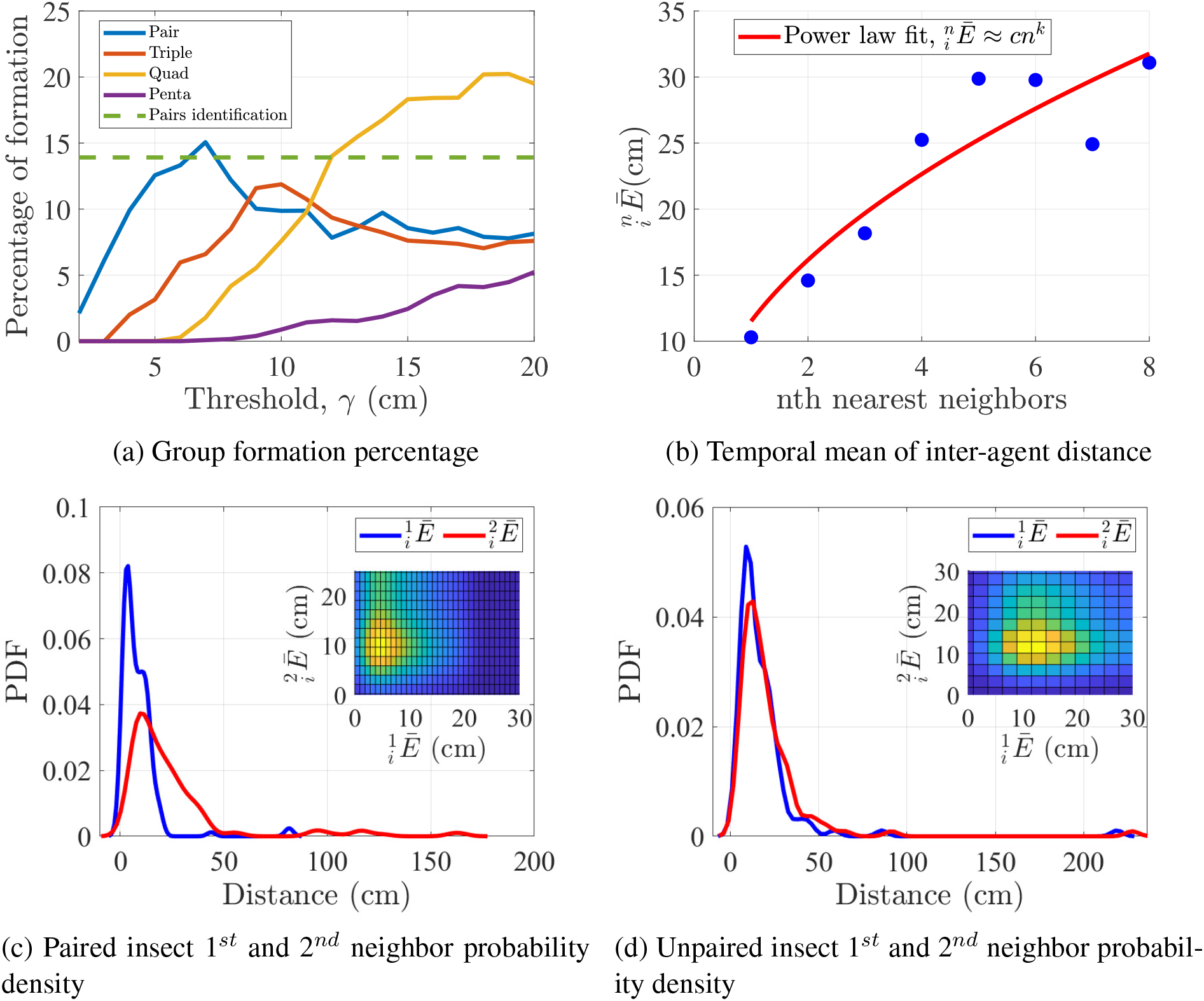
The percentage of group formation as a function of threshold (a) reaches a peak of 16% paired insects at *γ* = 7 cm, consistent with nearest neighbor analysis. The temporally averaged inter-agent distance follows a power law (c) *cn*^*k*^ by the *n*th nearest neighbor, allowing one to compare 1^*st*^ and 2^*nd*^ nearest neighbor’s distance (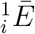 and 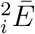 respectively) joint probability density functions (PDF), revealing that paired insects (c) show a significant distance between nearest (blue) and second-nearest (red) neighbors while unpaired insects (d) show a weaker separation.

To avoid sensitivity to the method used in neighborhood analysis, a third distance-based approach to neighborhood identification was incorporated (Refer to Methods: Pair bonded insects identification). This method used the entire trajectory of each agent and calculated their average neighbor distance 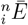, where *n* is the neighbor from agents set {1…*N* − 1}. The temporal average of distance 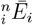 of *n*th nearest neighbors followed a power law prediction, as seen in Fig. 4b.

The power law exponent was *k* = 0.49, indicating that the first neighbor distance 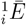 and second neighbor distance 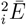 of a pair of birds must satisfy the condition 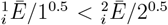. On the other hand, non paired birds satisfy 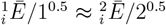. Figure. 4c illustrates that in paired birds, the PDF of 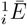 has consistently remained low, even as the PDF of 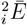 has shown an increase which suggests a decrease in local density. Conversely, non-paired insects exhibit very similar PDFs, as shown in Fig. 4d, indicating that 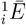 changed proportionally with 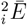. Thus, we can consider two insects *i, j* as being paired if their temporal average inter-agent distance is smaller than 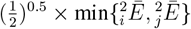 (refer to Methods: Pair bonded insects identification). This pairing method and the threshold, *γ* = 7.0 cm gave almost similar percentages of paired insects in the data which is around 14% as seen in Fig. 4a indicating on average 28% insects made pair formation while flying together.

### Pairwise regulation analysis

In these paired flights, the insects typically demonstrated pairwise tracking, as seen in S2 Video. These will be referred to as leader-follower behaviors by labeling the insect nearest the moving stimulus as the leader and the insect approaching it as the follower. Visual signals are the dominant signal modality available in-flight to support close-range coordination, raising the question *Does the follower adjust (regulate) its flight path to achieve this pairwise movement and if so, to what signals does it respond?* Hypothesis **(H1)** is that during pairwise flight, the rear (follower) bee adjusts its flight based on measurable visual signals of the leader. To test the visual response hypothesis, this study quantified three visual signal candidates: optical expansion rate *R*_*o*_(*t*), optical flow *R*_*f*_ (*t*) or relative velocity *R*_*v*_(*t*) (see Methods: Regulation candidates) that could be used to regulate relative to other agents or moving visual targets. To test the possible hypotheses, we quantified the three regulation candidates throughout the group behavior. Figure 6a shows the root mean squared error (RMSE) values for the three candidates, indicating the existence of three zones. In the “locked” approach zone the RMSE error for the optical expansion rate *R*_*o*_(*t*) is lowest and the two other candidates *R*_*f*_ (*t*) and *R*_*v*_(*t*) showed higher error (RMSE) (see Methods: Regulation candidates). The decision zone shows RMS error is similarly high for all three candidates as seen in Fig. 6b. Lastly, the response to the *R*_*v*_(*t*) candidate is best observed in the third velocity matching zone, as depicted in Fig. 6c.

Fig. 7 shows the followers’ average velocity, the leader-follower inter-agent distance, and the optical expansion rate as seen by the follower. Fig. 7a shows that followers maintained a constant optical expansion rate with less than 8.7% variance beyond 10 cm inter-agent distance. An example of a follower insect and its velocity and optical expansion rate *R*_*o*_(*t*) are shown in Fig. 5 and supplementary video S3 Video. Figures 5a and 5b show their 2D and 3D positions. The follower maintains a regulated optical expansion rate *R*_*o*_(*t*) in Fig. 5c until inter-agent distance reduces to 10 cm. As the distance decreases below 10 cm, *R*_*o*_(*t*) rises. The insect remained in this un-regulated region for 0.60 seconds and then re-engaged the leader with velocity matching. *Could the presence of an unregulated period between differing visual regulation behaviors be induced by the presence of a moving hive stimulus or would it persist when the hive is static? Similarly, would the three zone behavior persist if insect pairs did not respond to a stimulus in view?* To determine whether the behavior was an artifact of the artificial moving stimulus, the experiment was repeated with the moving stimulus held constant. The 3-zone behavior persisted with the frozen hive stimulus, and an example is shown in supplementary video S6 Video. As demonstrated in the supplemental video S7 Video, the behavior continued even in response to a moving stimulus. In this frozen stimulus example, the interacting insects did not follow the frozen stimulus, as indicated by their low correlation coefficient of 0.32, suggesting insect interaction was stronger than stimulus interaction.

**Figure 5:**
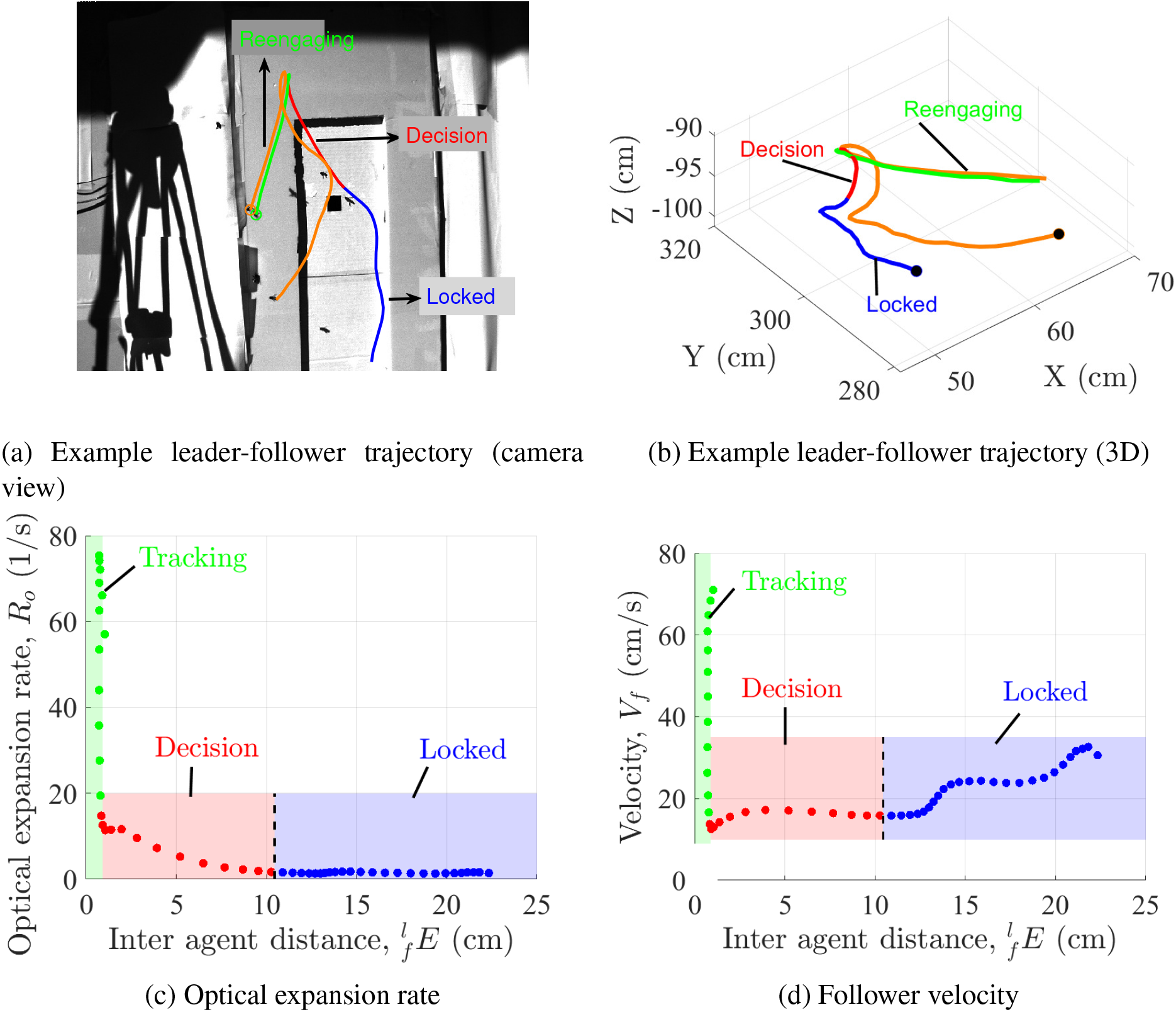
An example leader-follower trajectory (a) shows three behavioral zones, indicated with follower trajectory color (blue, red, and green); which are also visible in 3D trajectories with initial positions indicated in black. Optical expansion rate *R*_*o*_(*t*) in (c) obtained using Eqn. 7 illustrates the low variability during the locked region and subsequent lack of regulation, while follower velocity ***V***_*f*_ (*t*) in (d) lack a similar tightly regulated region.

**Figure 6:**
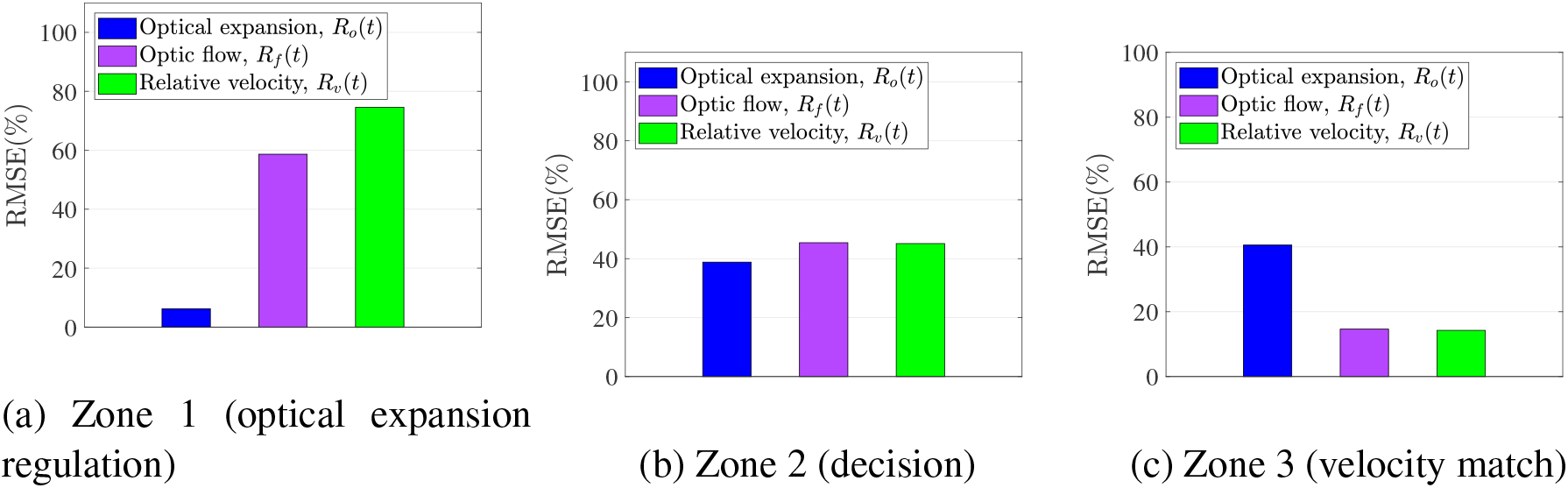
Root mean square error (RMSE) for three candidate regulation parameters indicates that follower insects initially regulate optical expansion rate (zone 1), then cease regulation during observation and decision (zone 2). Those agents that re-engage after zone 2 then match the leader agent’s velocity (zone 3).

**Figure 7:**
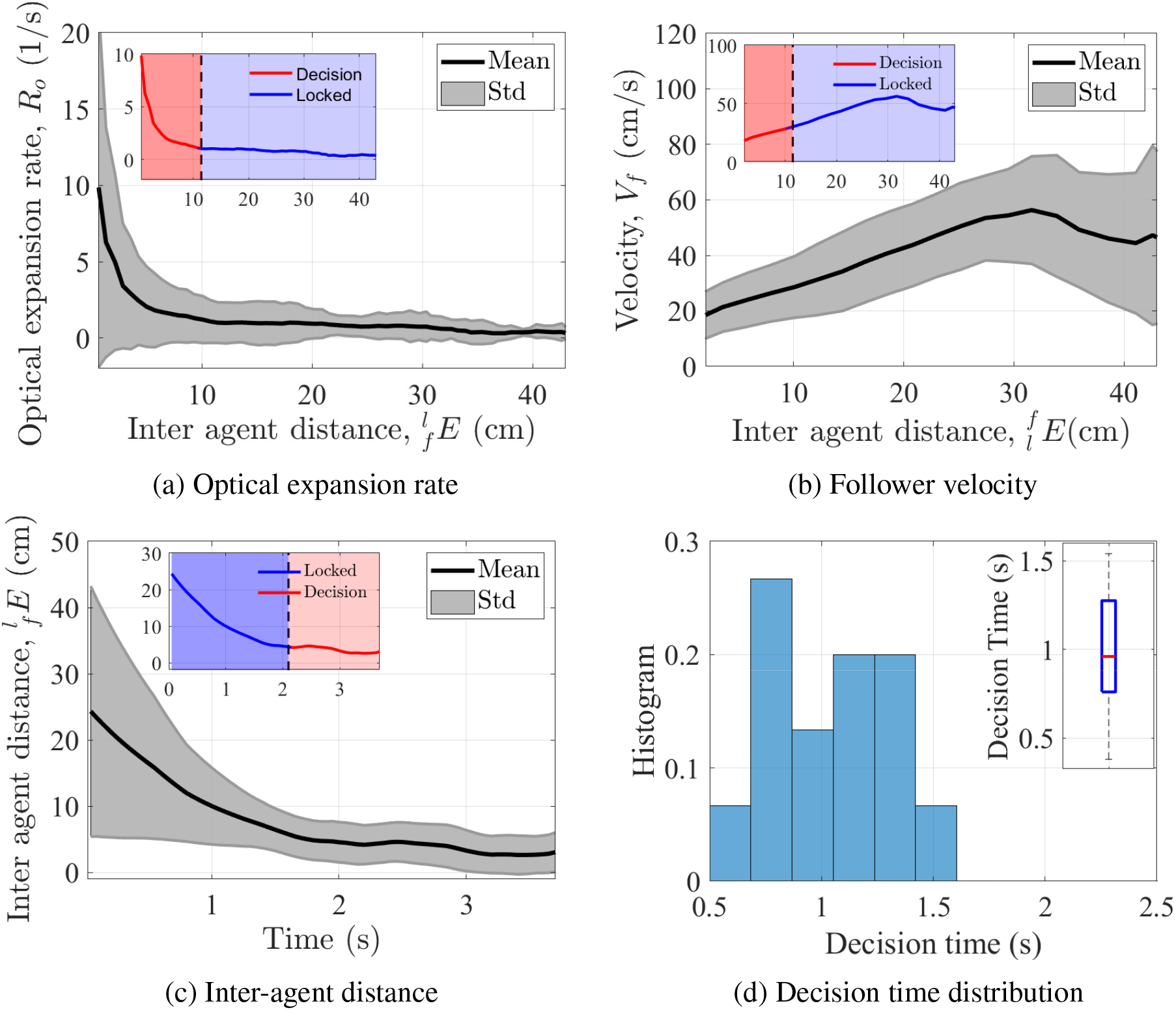
Optical expansion rate (a; showing mean in black and standard deviation in grey) was regulated with 8.6% variance beyond 10 cm inter-agent distance. Optical expansion rate is composed of follower velocity (b) inversely scaled by inter-agent distance (c), whose largely inverse relationships with respect to distance showed the effect of regulation. The distribution of time spent in the observation/decision zone (d) varied from 0.5 to 1.5 seconds, with a median of 0.97 seconds.

The horizontal plane heading of the two agents was analyzed via a position-only heading analysis and a velocity/position analysis as described in Sec. Heading angle analysis. The orientation of a line connecting leader and follower positions showed higher variations beyond 10 cm inter-agent distance than below 10 cm. The orientation of the follower’s velocity relative to this leader-follower position line also showed low temporal variation (as quantified by its time derivative) for paired insects nearer than 10 cm than beyond 10 cm, where its time derivative showed high variability. Together, these two outcomes–and their existence from the lateral-directional domain–reinforce the existence of a 10 cm metric distance trigger between behaviors.

### The un-regulated observation decision zone (zone 2)

After the inter-agent distance had closed to 10 cm, the follower entered an intermediate region that would eventually lead to an engagement or disengaing behavior. The length of this intermediate region motivated the study to consider *how could this intermediate zone support the ensuing engagement/disengagement decision at the conclusion of the zone?* Hypothesis **(H2)** is that follower insects implement a range-specific entry to an in-flight observation and deliberation zone to engage/disengage the pairwise leader.

In Fig. 5b, the three identified zones–locked, decision, and reengagment–are delineated with varying colors. Fig. 7d shows that the overall time in the decision zone is variable, with a median time of 0.97 seconds.

Insects spent an median of 0.97 seconds deciding to reengage or disengage, and in 79% of cases the followers followed the decision-making zone with velocity tracking. Several disengagement decisions are shown in movie S4 Video. The existence of deliberative observation and decision-making in insect flight is supported by this discovery of a long temporal region in which the visual candidates cease to be regulated, and that the animals re-engaged the same leader in 79% of the trials and disengage in 21% of the trials.

### Heading analysis

Analysis of the relative heading from leader-follower trajectories was conducted to figure out any specific patterns in heading angles. Two different approaches were employed. Initially, utilizing the relative angle method (detailed in Method: Position-based heading analysis), the heading angle from followers to leaders was computed, exemplified in Fig. 15a. Subsequently, the heading angles of all leader-follower pairs were processed in batches, as illustrated in Figure 15b. The heading angles consistently decreased from the approach zone to the decision zone, indicating a general downward trend. However, this observation provides limited additional insight beyond the overall reduction in heading angles. For a more comprehensive analysis, an alternative method based on a force map to find the relative change of angle and speed between pairs of insects was implemented (refer to Method: Position/velocity-based heading analysis) (Mudaliar and Schaerf, 2020). The average speed and angle change of the followers with respect to leaders were visualized in the relative coordinate system’s force mapping. A detailed method for generating the map of the average change in speed and angle is provided in the Method section (refer to Method: Force map). This force map illustrates followers’ tendencies to orient or adjust speed within pairwise movement, detailing how followers modulate their direction of motion and speed relative to leaders as a distance function. The individual probability density *J*(*x*, 0), *J*(0, *y*) and joint probability density *J*(*x, y*) of the follower’s *x* and *y* coordinates in the relative force map are shown in Fig. 16. The absolute average speed change of followers in relative coordinates is illustrated in Figs. 17a and 17b which demonstrates that followers experience less variability in speed changes when they are closer to leaders compared to when they are farther away, a threshold value of 70 can be used to distinguish the lower variability region (red shading) seen in Figs. 17a and 17b. Specifically, for the *x* coordinate, the variability ranges from −7 to 10.4 cm, and for the *y* coordinate, it ranges from −7 to 5.7 cm. The 3D representation of this speed change is provided in Fig. 18b, with the lower speed change depicted in blue. The changes in followers’ angles are also represented in Figs. 17c, 17d and 18c, which reflect similar trends observed in speed changes, with lower angle changes (variability *<* 3.5°) indicated by red color near the leaders, with ranges of −5.1 to 8.7 cm for the *x* coordinate and −4.9 to 4.1 cm for the *y* coordinate. The reduced variability at closer distances suggests that the followers might not try to match the speed or angle, indicating a disregard for the regulation in favor of making independent decisions. Conversely, at larger distances, the followers adjusted their angle and speed to align with the leader. Additionally, the relative direction of followers can be used to assess group orientation alignment. Figure 18a illustrates the average relative direction of follower motions where the arrows denote the alignment, with straighter arrows indicating a higher degree of directional match between leaders and followers.

### Re-engagement regulation (zone 3)

When re-engagement was achieved, the first question is *by what rule do the follower insects regulate their velocity with the leaders in the re-engaing zone?* The RMSE associated with velocity and optic flow initially appear similar in Fig. 6, however the mean error difference of 0.39 exceeds the variance difference of 0.87, and standard deviation difference of 0.0882, indicating that the velocity RMSE is significantly lower, and the experimental data better supports a velocity regulation explanation than an optic flow regulation interpretation. This analysis then considers a PID feedback structure to govern paired movements wherein variable integral gains may serve as an indication of short timescale adaptation **(H3)**. The insect trajectories where the follower re-engaged its leader were considered to determine the regulation strategy used in the velocity matching zone. As the stimulus moves along the *ĝ*_*y*_ axis of the world frame, the dominant movement of the insects is reflected by this motion and was used to determine the feedback rule. A PID feedback control structure (shown in Fig. 8a) was used to compute feedback gains *k*_*p*_, *k*_*i*_, and *k*_*d*_ from the leader and follower velocities (refer to Method: Feedback control rule (PID)). S5 Video shows an example of two insects’ paired rengaging behavior and their 3D positions are shown in Fig. 8b, illustrating the dominant movement in *Y* axis. The leader velocity ***V***_*l*_(*t*) and follower velocity ***V***_*f*_ (*t*) are shown in Fig. 8c. For this example in Fig. 8d, the *k*_*p*_, *k*_*i*_, and *k*_*d*_ gains are 18.5594, 1.1012, and 0.0296, respectively, and the FIT percentage according to Eqn. 32 is 76%. Across the dataset, Figures 9a and 9b show the proportional and integral gains *k*_*p*_ and *k*_*i*_ versus the average inter-agent distance 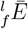. The integral gains vary while the proportional gains have a median value of 23.24 as seen in Fig. 9c. The relation of the two gains is shown in Fig. 12a. Figure. 12b shows that the identified *k*_*d*_ gains are always less than 1 and negligible compared to *k*_*p*_ and *k*_*i*_, thus the velocity matching behavior in zone 3 is best modeled with a PI control structure.

**Figure 8:**
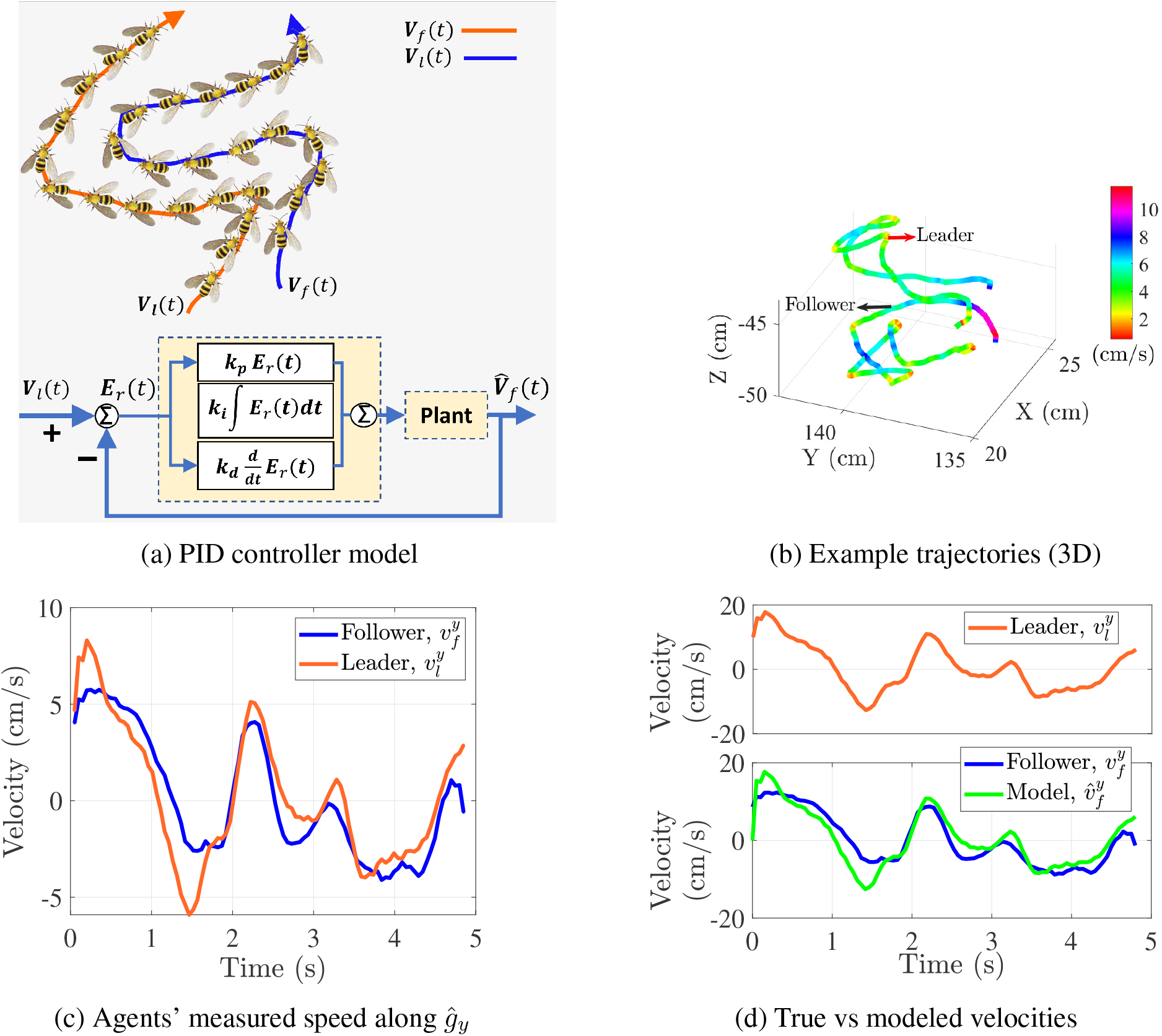
A PID controller model (a) was considered to examine follower’s velocity 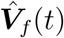 in zone 3 in terms of leader velocity ***V***_*l*_(*t*) as a reference input. Example zone 3 leader and follower trajectories are shown in (b) with color indicating instantaneous overall speed |***V***_*i*_(*t*)|, while considering only the measured vertical axis speed reveals a velocity-matching behavior. This behavior is well-described by the PID control structure in (a) and in Eqn. (28-30), as verified via the the simulated model velocity 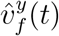 shown in (d) populated with identified PI gains.

**Figure 9:**
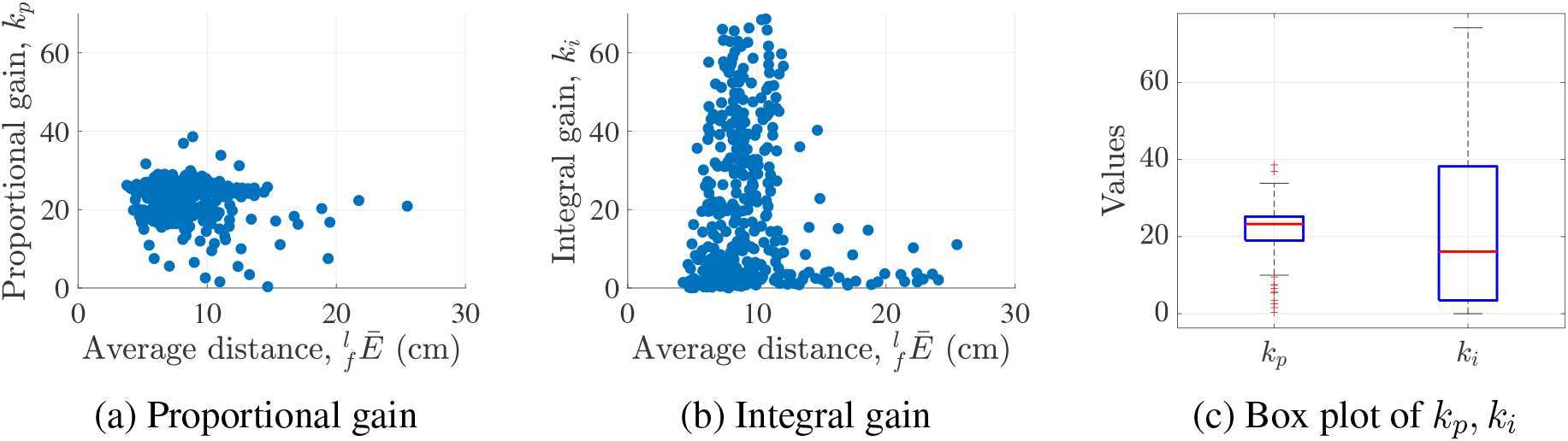
The inter-agent specific variation of proportional and integral gains *k*_*p*_ and *k*_*i*_ revealed a relative consistency in proportional gain (a) and a variability in integral gain (b), which is also visible in the gains’ interquartile and median values in (c).

## Discussion

In this study, we examined the behaviors of 1000 honeybee group flights as they approached the entrance of a beehive. Our aim was to understand the formation of group flights and explore how their in-flight state variables were modulated to regulate pairwise movement.

### Honeybees exhibit pairwise flights near hive entrance

Anisotropy analysis indicates flocking starlings interact with six to seven topological neighbors, while insects’ limited neural and optical systems suggest they may use a smaller interaction range. Cross-correlation analysis between neighbors of a given honeybee, akin to determining the correlation length or spatial span, indicates 2-4 interacting neighbors, as illustrated in Fig. 3. The direct interaction range could be even more limited (Cavagna et al., 2010) and a cross-correlation finding does not fully negate a metric distance explanation. Inter-agent distance analysis shows a threshold sensitivity to identifying group size, and indicated up to 32% insects participate in pairwise flights, which would be achieved when considering a geometric group size of 7 cm. Pairwise flights were also identified by considering the relative distance between 1st and 2nd nearest neighbors, indicating 28% of agents particiated in pairwise flights as seen in Fig. 4. These paired relationships became the basis for the feedback regulation analysis. Animals in coordinated flight typically interact with a limited instantaneous number of neighbors, and three methods for neighborhood analysis in the current study consistently indicated approximately 30% of involved agents show interactions with only a single neighbor. This interaction neighborhood is smaller than previously quantified birds, potentially as an adaptation to insects’ more limited optical and neural resources. The pairwise coupling may still support coherent motion coordination through feedback regulation.

### Honeybees initially regulate the optical expansion rate in pairwise interactions

Previous studies have examined constant bearing angle, absolute target direction, and multisensory motor rules in animal pursuit behavior. These predator/prey and mating pursuit stategies do not describe the partners in peer-relationships, such as the interactions in high density forager-forager traffic, where this study showed a the domimant interaction is pairwise. To explore a possible strategy when inflight neighbors interact with each other, an output regulation hypothesis (H1) was tested in the outdoor group flying experiment by comparing the error of a regulated value. This analysis showed that of the three candidates (optical expansion rate, optic flow, and relative velocity), honeybees regulated the target’s optical expansion rate when they followed a leader at a relative distance greater than 10 cm. The optical expansion rate begins to increase at an inter-agent distance of less than 10 cm as depicted in Fig. 5. Heading angle analysis similarly revealed a critical distance of 10 cm in the paired flights, beyond which followers modulated their angle and speed to maintain a constant optical expansion rate, resulting in spikes and fluctuations as seen in Fig. 18. Once inter-agent distance had reduced to 10 cm, these adjustments ceased, suggesting the absence of a fixed rule governing behavior in this region. In light of these findings, the approach region from acquisition to 10 cm closure was labeled zone 1, the “optical lock” zone.

Optical expansion rate has been quantified in previous solitary studies, such as its use as a landing cue or approach signal (Goyal et al., 2022; Reiser and Dickinson, 2010). However, in landing and solitary navigation studies, the visual target, such as a surface to land on, is stationary. In the context of this study, the follower insect regulates *R*_*o*_(*t*), a term dependent on its own absolute velocity scaled by the relative distance to the leader (Eqn. 7). This fraction of relative and absolute quantities is remarkable, and distinguishes the quantity from the optical flow signal *R*_*f*_ (*t*) that incorporates relative quantities and has considerable support as a basis for in-flight navigation (Mauss and Borst, 2020; Serres and Ruffier, 2017; Srinivasan and Zhang, 2000). One interpretation for using *R*_*o*_(*t*) instead of *R*_*f*_ (*t*) is that the insect lacks an ability to estimate the leader’s velocity, either due to the limited angular size and thus stimulus, or because of limitations in neural processing needed to estimate another agent’s velocity. The terms are equivalent when the observed target (leader) has zero velocity, suggesting the possibility that a similar neuromusculature process governs both. Insects in solitary flight do have an ability to tolerate behavior-gated differing levels of optical expansion (Reiser and Dickinson, 2010), demonstrating an interplay between navigaiton and expansion tolerance, thus determining whether the the neural processing chain active in this crowded approach behavior incorporates an assumption of zero target motion would require a specialized study.

### In-flight observation and decision-making zone

Previous work has explored insect colony nest site decision-making and individual forage tracking and landing decisions. However, it is not clear whether the flying insects incorporate deliberative aerial decision-making in the sense of incorporating a lengthy observation period before initiating a binary decision, particularly in regards to which or whether to regulate relative to a neighbor. The current study’s finding that the same leader is consistently re-acquired upon re-engagement supports interpretations that either that an observation continues during the unregulated region or that the leader’s post-regulation stimulus is distinct enough to support reacquisition. No clear difference in leader trajectories to support a re-acquisition was observed in this study, suggesting the observation/decision hypothesis may be stronger. This individual decision-making is distinct from colony-level decision making such as honeybee nest selection in which a minority of individuals facilitates a group decision.

This study explored the possibility that honeybees are able to make an in-fight decision about whether to re-engage or abandon pursuit of a conspecific they are tracking (Hypothesis H2). In pairwise leader-follower dynamics, the analysis indicates that followers exhibit a measurable deliberative period initiated at a 10 cm inter-agent distance. None of the visual regulation candidates were regulated after the 10 cm radius initiated zone 2, as seen in Fig. 6. The duration of these decision-making processes varied across the dataset (Fig. 7d), with a median decision time of 0.97 seconds. The length of this observation and decision process is significantly longer than previous in-flight decision measurements (Wardill et al., 2017; Lin and Leonardo, 2017; Talley et al., 2023). The significant variability in decision time suggests that the decision-making process during flight is not uniform and may be adaptable across individual or environment.

#### Follower heading angle adjustments in pursuit strategies

In the pairwise analysis, the relative heading angle between insect pairs exhibited a decreasing trend (seen in Fig. 15b). Additionally, the follower’s change in angle and speed relative to the leader showed greater variation at larger distances and smaller variation at closer distances as seen in Fig. 17. High and low variation thus segments the behavior spatially into two regions: Zone 1 and Zone 2. In Zone 1, the follower pursues the leader, while in Zone 2 less change is seen, in consistent with the observation/decision zone seen earlier. These zones may represent two different strategies: a pursuit strategy in Zone 1 and a decision-making strategy in Zone 2. An analogous distinction has been found blowflies, which identified two different pursuit behaviors—one for target tracking in the horizontal plane and another for target interception in the vertical plane Varennes et al. (2020b). The difference arises from the angle of error between the body axis and the speed vector. In the horizontal plane, this angle is close to zero, allowing for more accurate tracking. However, in the vertical plane, there is a misalignment between the speed vector and the line of sight, leading to differences in pursuit behavior. In this study, followers may employ two distinct approaches in their pursuit strategy when following leaders. In Zone 1, followers exhibit high variation in angle and speed changes to closely track the target. In contrast, in Zone 2, they relax these variations, resulting in more stable movement, as shown in Fig. 17.

Pursuing *Drosophila melanogaster* attempt to stabilize head position under slow rotating polarized light, though heading movement varies across individuals Warren et al. (2018). For example, three different *Drosophila* exposed to the same rotating UV light showed mean heading angles of 140°, 107°, and 115°, with standard deviations of 10°, 27°, and 13°, respectively. Another study revealed similarly stable headings relative to the angle of polarized light, but the heading distribution was broad, with only a slight deviation from a uniform distribution Mathejczyk and Wernet (2019). This individual variation in *Drosophila* individual heading during the pursuit strategy reinforces the observed high variation in Zone 1 in this study.

### Follower insects use PI feedback rules to match coupling flights

This study found that across 1000 trials, 79% of paired insect recordings showed follower reengagement, and the re-engaged feedback rule fit a PI control structure as illustrated in Fig. 8, with consistent proportional gains (Fig. 9a), variable integral gains (Fig. 9b), and negligible derivative gains (Fig. 12b). A proportional integral structure has previously been observed in *Drosophila melanogaster* solo flight the *b*_1_ and *b*_2_ wing muscles’ role in stabilizing pitch motion (Whitehead et al., 2022), suggesting a muscle function specialization each muscle in flight control. The current study finds that the inflight PI construction seen in pitch stabilization may be a generalizable architecture integrated into more complex behaviors more complex more diverse behaviors, such as neighbor-relative flight control in groups. Predator-prey tracking, such as blowfly experiments, have concentrated on primarily reactive (proportional) feedback such as pure-pursuit, biased pursuit, constant bearing angle proportional navigation (Varennes et al., 2020a). A proportional-only controller performs well on integrating processes not requiring a biased input, while PI control includes a “moving bias” and more applicable for “non-integrating processes,” or those that would otherwise eventually return to the same output given the same set of inputs and disturbances. The short term adaptation in PI control can act as a storage term, or a short term memory, that results in longer term energy trade-offs visible as oscillations. The finding that the leader-follower tracking structure in flying groups includes an integration beyond the proportional rule found in prey pursuit suggests that conspecific group interactions may encode fundamentally different feedback approaches, and may reflect differing task structures. This knowledge of the structure can guide extensions, such as the inclusion of controllers using the internal model principle.

### Reactive, moderate, and deliberative timescales

Human psychology analysis incorporates multiple timescales, most popularized by Kahneman (2011), which modeled human information processing by two concurrent systems that exhibit transient dominance. A “fast thinking” (System 1) operates automatically and intuitively, while a ‘slow thinking” (System 2) involves deliberate, focused, and concentrated processing. The current study broadens this approach by recognizing that the insect behavior includes a sequence with a fast system (reactive control in region 1, typically 7-16ms loop closure rates (Autrum, 1958; Vance et al., 2013; Ristroph et al., 2013)), a slower deliberative process that spends a median of 0.97 second in the decision region 2), and a medium timescale (PI control in region 3) in which reactive behaviors are combined with a slower mathematical integration to produce a moderate timescale behavior (Tavares, 1966). The finding of three subsequent behaviors that show significantly different timescales raises the possibility that insect systems may support processing on a spectrum of timescales beyond the “fast” and “slow” systems previously identified, a promising area as autonomy seeks to implement more diverse constructions.

### Alternative swarm models

Swarm constructions often incorporate long range attraction, mid-range velocity alignment, and short range repulsion. Control theoretic paradigms often leverage an engineered separation of guidance, navigation, and control (GNC) disciplines, while robotic approaches involve sensing, perception, decision, and action (SPDA) paradigms (“sense, decide, engage” or “observe, orient, decide, act” in engineered military contexts) often mirroring the same three functions of velocity alignment, attraction, and collision avoidance (Reynolds, 1987; Cucker and Smale, 2007). Agents in biological groups incorporate more individually diverse constructions. The “lock on/approach, decide, then track” behavior discovered in this study and summarized in Fig. 10 is an alternative construction for coordinated groups. The architecture may be an example of a construction that could reduce swarming’s computational and attentional requirements (Billah and Faruque, 2024) and accommodate more diverse embodiments, such as those with non-Turing processors, limited communications, lack of absolute position references, and a desire for apparently less structured motions.

**Figure 10:**
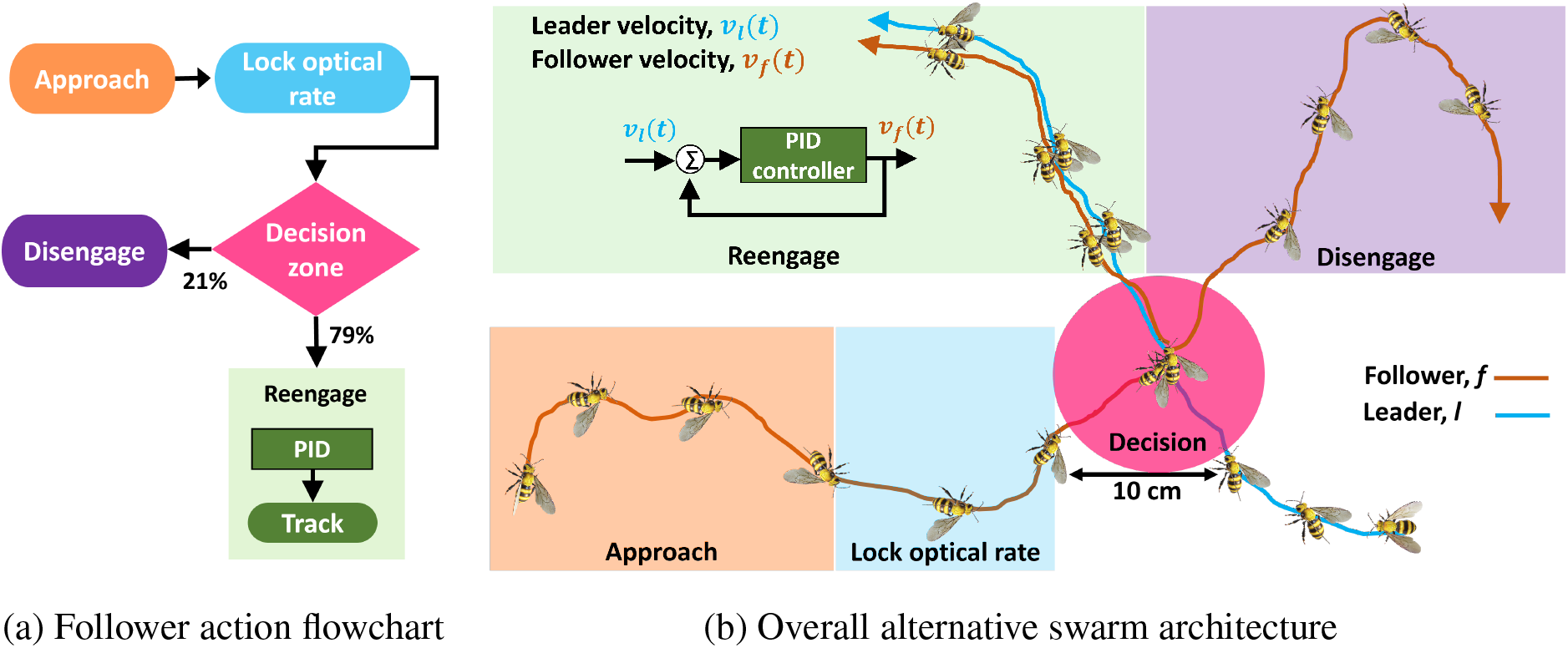
The follower’s actions towards the leader are summarized in a flowchart (a), and the overall leader-follower relationship shown in (b), which highlights the follower insect’s progression: approach, lock optical rate, decision zone (entry triggered at 10 cm), and either PI velocity match or disengage.

## Conclusion

The study identified the interaction network that supports visual coordination in honeybees in crowded hive entrance approach regions. This study first quantify the interaction range of honeybees flying in crowded hive approach regions, then isolated the feedback and decision-making architectures demonstrated throughout this group interaction. The interaction range analysis reveals that honeybees primarily interact with one to four other agents within their visual range, with the largest fraction of insects engaging in pairwise interactions. This network’s strong reliance on pairwise interactions allowed identifies the pairwise decision times and changing feedback control rules in flying insects. The study revealed that an individual bee regulates optical expansion rate, enters an observation/decision region, and then either re-engages with the leader in proportional-integral (PI) velocity error or disengages from group behavior. The analysis highlighted the relatively large fraction of time spent in decision-making during the dynamic interaction. Together, these new results uncover an alternative visual swarm model summarized in Fig. 10, that incorporates optical feedback, low network (connectivity) requirements, and integrated-decision making.

The coordination supported by this behavioral combination suggests an alternative to the traditional swarm and flocking motion models that often focus on achieving consensus in motion variables like heading and velocity rather than incorporating dynamic individual agents’ decision-making abilities in motion prediction.

This alternative model’s reliance and low connectivity and optical signals is also a foundation for high speed swarming aerial robotics with limited onboard computational resources. Understanding how animals reduce the attentional requirements to support stable motions can aid engineers in allocating processing tasks effectively in such large scale partially observable systems, especially by reducing the overall network traffic requirements. This integrated architecture holds promise for diverse applications, including search and rescue, environmental monitoring, and surveillance.

## Materials and Methods

### Imaging system

The experimental apparatus is divided into two parts: visual stimulus and tracking system. The stimulus is a fixed or vertically moving beehive entrance (rectangular hole of 5 cm). The stimulus was moved vertically using a combination of sinusoidal frequencies through two stepper motors. The image processing software VISIONS captured and quantified the three-dimensional positions of the insects (Islam and Faruque, 2022). The system requires three or more cameras placed in front of the flying arena. The camera frames were recorded at 50-120 frames/s and 640 × 512-pixel resolution. Three scenarios were considered: (a) the follower and leader approaching the moving stimulus, (b) a static stimulus, and (c) and the leader and follower flying away from the stimulus.

### Insect stocks and handling

At Oklahoma State University, experiments were conducted using a hive of western honey bees (*Apis mellifera*). The honeybees’ flight trajectories were recorded during between 12 pm and 4 pm within a flight arena measuring 150×150×150 cm. The stimulus and tracking system were operated remotely to minimize disruption to their natural behavior. Group flights typically consisted of three to ten insects.

### Correlation range

To find the information transfer in a group of flying insects, this analysis first considered a pairwise-directed graph network system. When an agent *i* interacts with other agents *j* it may send or receive information from them. If the information transfers among themselves, an adjacency matrix of the followers and influencers can be established. The graph of the network will have a set of vertices (nodes), *V* = {1, …*N*}, indexed by the *N* agents of the group. The set of edges, *ϵ* = {(*i, j*) ∈ *V* × *V*} contains the pairwise connections in the group. As an example, let us consider four insects flying toward the stimulus. If any agents receive information from others, the corresponding weight will be non-zero. In Fig. 3a, agent 2 receives information from two other agents (agent 3 and agent 1), while agent-4 receives information from all three other agents. The adjacency matrix ***W*** of this example is

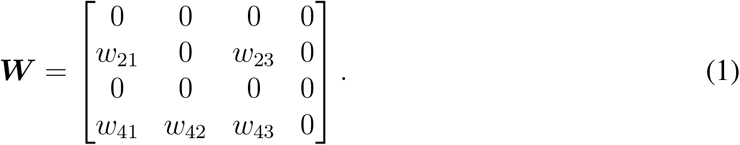

One method to estimate weights *w*_*i,j*_ is cross-correlation analysis. The inter-insect cross-correlation is defined by considering a time delay *τ*. Let *y*_*i*_(*t*) and *y*_*j*_(*t*) be the *y* coordinates of insects *i* and *j* and 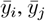 be the mean of *y*_*i*_(*t*) and *y*_*j*_(*t*), respectively. Then, the cross-correlation *S*_*i,j*_(*τ*) is

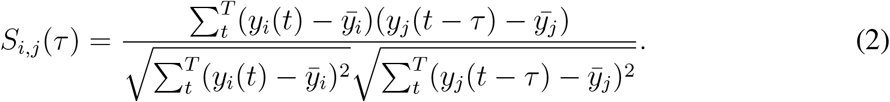

Cross-correlation analysis varies the lagtime as *τ* = [− *T, T*], and finds the maximum value of *S*_*i,j*_(*τ*), where *T* is the total time length. When agent *i*’s action precedes agent *j*’s similar action, cross-correlation analysis interprets agent *j* as receiving information from agent *i*. The maximum correlated magnitude 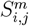 can be expressed as

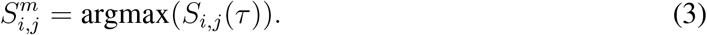

To form a directed network, the weight *w*_*i,j*_ of two agents is determined by

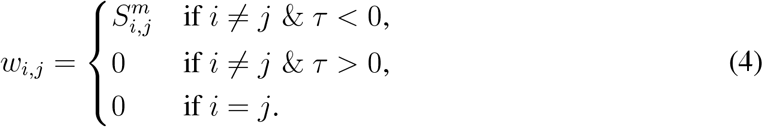

The direction of information flow will depend on the sign of *w*_*i,j*_. If *w*_*i,j*_ is positive, agent *i* receives information from agent *j*. Conversely, if *w*_*i,j*_ is negative, the direction of information flow is reversed, meaning agent *j* receives information from agent *i*. An example of six insects’ 2D and 3D positions is shown in Figs. (3b-3c). The *y* coordinate of each insect shown in Fig. 3d is used to find the adjacency matrix among themselves. This analysis considered the *y* coordinates because the stimulus moves along the world *ĝ*_*y*_ axis. Cross-correlation analysis estimated the adjacency matrix for this example as

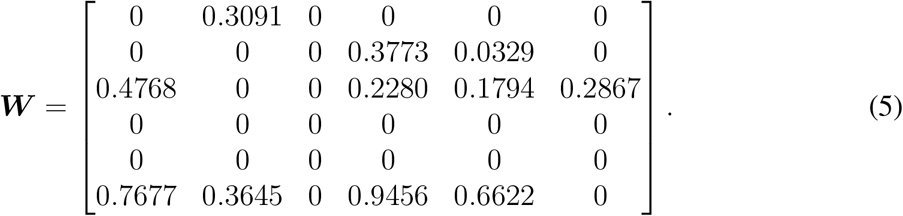

An agent receiving information (in-degree) is indicated by a non-zero value. One can find the Laplacian matrix ***L*** from the adjacency matrix ***W*** by ***L*** = ***M* − *W***, where ***M*** is the degree matrix of the graph (Mesbahi and Egerstedt, 2010; Bullo, 2024). The second eigen-value (Fiedler number) of the Laplacian matrix is 0.869 which indicates the graph has multiple connected components. If the Laplacian matrix has a single eigenvalue equal to zero and all other eigenvalues have positive real parts, the graph is strongly connected and has at least one spanning tree (Mesbahi and Egerstedt, 2010; Bullo, 2024). The above example is a strongly connected graph, shown in Fig. 3e. This study also considered the graph’s matrix norm, degree distribution, in-degree /out-degree weight statistics, as seen in Table 1 (reporting the norm and in/out degree weight medians) and Table 2 (showing the the in-degree connectivity values) of this adjacency matrix ***W***.

**Table 1:** Matrix size and median weights.

| Properties | Value |
| --- | --- |
| Norm ($\vert \vert \mathbf{W}\vert \vert $) | 1.543 |
| In-degree median weight | 0.771 |
| Out-degree median weight | 0.772 |

**Table 2:** Connectivity statistics.

| Number of connectivities | Number of insects | Fraction of insects |
| --- | --- | --- |
| 1 | 1 | 0.16 |
| 2 | 1 | 0.16 |
| 4 | 2 | 0.33 |

To analyze the set of multiple trials, this study first concatenated datasets vertically and applied the cross-correlation analysis technique. This norm of the matrix increased with the group size, indicates that as the number of insects increases, their mutual communication also increases, as shown in Fig. 3f. The summation of average indegree and outdegree weights for each node of the group of flying insects in Figs. 3g and 11 respectively, indicates that as the size of the insect group increases, their average indegree and outdegree weight sums also increase. Consequently, the average amount of transferred and received information per insect rises with group size. This analysis indicates most (70% in Fig. 3) of the insects are cross-correlated with 2-4 neighbors regardless of their inter-agent distance in Fig. 3h.

### Identification of nearest neighbors

#### Group sizes sorted by distance threshold

A inter-agent distance threshold based method was also used to find the number of insects in each group. If the average temporal inter-agent distance along the entire trajectory 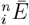 from a focal insect *i* to its *n*th neighbor falls under the threshold *γ*, one could consider those two insects (*i, n*) as an interacted insect pair. After this thresholding, a corresponding network graph is built for *N* agents and the weight of the adjacency matrix can be taken as

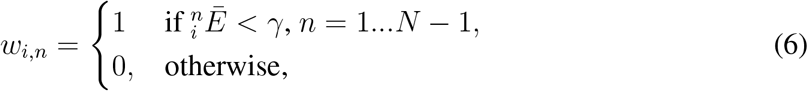

and the degree of the graph’s adjacency matrix reveals the number of insects in that group. In this way, We can sort the pair, triple, quad, and penta groupwise formations.

#### Pair bonded insects identification

Pair-bonded insects can be identified using the temporal average distance based method (Ling et al., 2019b). If the temporal average distance 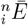 between a focal insect *i* and its *n*th nearestneighbor follows a power-law relationship 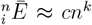, where *c* is a constant and the exponent *k* is approximately 0.5, then the method can be applied. For the overall insect trajectories, the temporal average distance follows a power-law pattern, as illustrated in Fig. 4b. Due to the presence of pairs, the first nearest neighbor distance 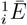 is lower than the power law prediction, paired insects must satisfy the condition 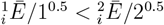, On the contrary, the unpaired insects exhibit 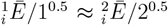. When these criteria are met, two insects (*i, j*) can be considered a pair if their temporal average 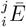 meets the condition, 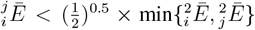 (Ling et al., 2019b).

### Regulation candidates

For convenience, each identified pair-bonded member was labeled as either a “leader” *l* or a “follower” *f*, with the member closest to the stimulus target labeled as the leader. Visual signals are strong cues in this interacting region of a leader-follower relationship, and this study quantified three visual candidates. The visual candidates considered are optical expansion rate, optic flow, and relative velocity. Given a follower velocity ***V***_*f*_ (*t*), leader velocity ***V***_*l*_(*t*), and the Euclidean distance 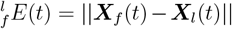 between their 3D positions ***X***_*l*_(*t*) and ***X***_*f*_ (*t*), the optical expansion rate *R*_*o*_(*t*) is defined as

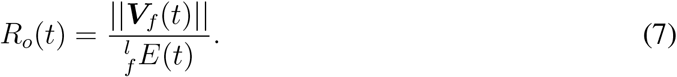

Similarly, optic flow *R*_*f*_ (*t*), and relative velocity *R*_*v*_(*t*) are

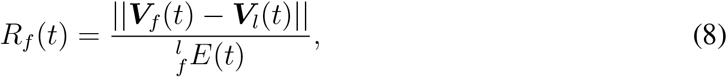

and

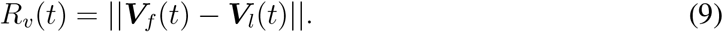

The analysis calculates the local slopes at each point of the visual candidate, which allows the identification of regions where they remain constant as the follower approaches the leader. To calculate the local slopes, we used finite differences. For example, if the optical expansion is *R*_*o*_(*t*) and the relative distance between the two insects is 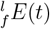, then the slope *p*(*t*) is the derivative of 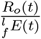 with respect to time. If the optical expansion rate remains constant, the slope of the (*R*(*t*) vs 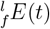) line will be close to zero, visible as in a horizontal in Fig. 7a. Consecutive points with slope *p*(*t*) change less than 8% were labeled as the “locked” zone.

This analysis showed that inter-agent approach distance greater than 10 cm is a locked zone in which the follower attempts to maintain a constant *R*_*o*_(*t*) relative to the leader. Optical expansion rate fixation ceased once inter-agent distance has reduced below 10 cm, and the other visual candidates also showed variation in this, suggesting an un-regulated region supporting observation and decision.

Our analysis considered this observation decision region to persist until the insects begin to either reengage or disengage. The duration from optical rate fixation cessation to re-engagement was labeled as the “decision time.” The time duration within the decision zone was measured for each trial. Once the follower starts reengaging, the relative velocity ∥***V***_*l*_(*t*) − ***V***_*f*_ (*t*)∥ begins to decrease, and this zone is labeled as the reengaging zone.

To identify which visual candidates the follower insects maintain in each zone (optical expansion regulation, decision, and velocity match), the root mean square error (RMSE) of *R*_*o*_(*t*), *R*_*f*_ (*t*), and *R*_*v*_(*t*) between leaders and followers was calculated as

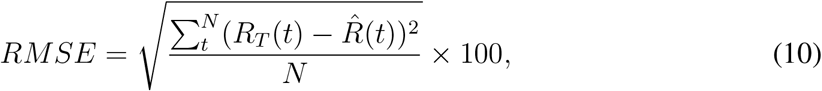

where set *R*_*T*_ (*t*) ∈ {*R*_*o*_(*t*), *R*_*f*_ (*t*), *R*_*v*_(*t*)} are the actual candidates and 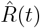 is the ideal case of the candidate. *N* is the number of data points of the candidates. If a follower regulated a candidate in any zone, the ideal 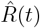 would be close to a fixed or constant value. Next, the overall trial-wise average RMSE values from all the leader-follower pairs were calculated.

### Heading angle analysis

Two approaches were used to analyze the headings of leader-follower pairs: a point-based and a relative motion-based methods. For heading analysis the *z* coordinate was discarded.

#### Position-based heading analysis

Considering two points ***X***_*l*_(*x*_*l*_, *y*_*l*_) is from a leader and point ***X***_*f*_ (*x*_*f*_, *y*_*f*_) is from a follower at a specific time instance in *G* frames as depicted in Fig. 14a. Then coordinates of leader can be calculated as,

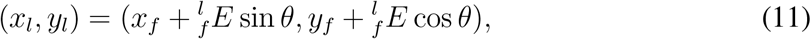

where 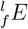 is the length from ***X***_*f*_ to ***X***_*l*_. It follows that *θ* satisfies the equation

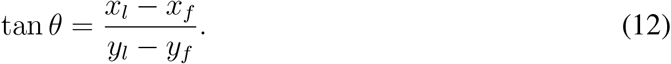

The calculation of *θ* using a four-quadrant arctangent (e.g., `>>arctan2`) function can be expressed as

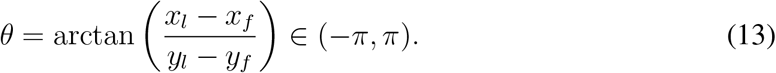

The relative heading angle *θ* ∈ (0, 2*π*) can be obtained by

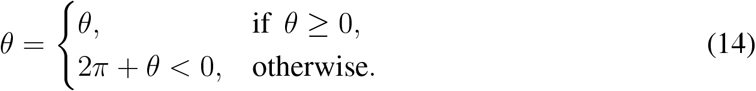

#### Position/velocity-based heading analysis

This method comprised two sections: firstly, analyzing the individual motion of the follower, and secondly, examining the relative motion between the follower and the leader (Mudaliar and Schaerf, 2020).

##### Individual motion

Initially, the velocity 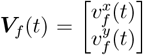 of the follower insect *f* at time *t* as seen in Fig. 14b are computed as follows:

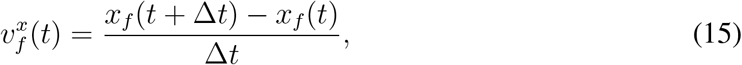

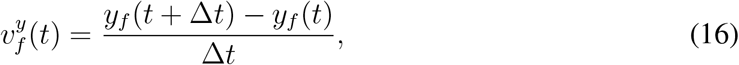

where, 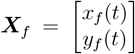 represents the follower *f* position at time *t*. The unit vector of the follower velocity is defined as

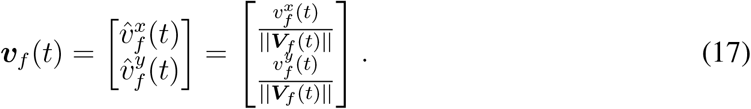

The magnitude of the direction of motion of the follower can be expressed as:

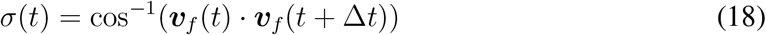

To determine the clockwise and anti-clockwise direction of the motion, the cross product of two unit vectors is used

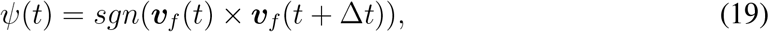

where × denotes the vector cross product. Thus, the signed change of the direction of motion is calculated as

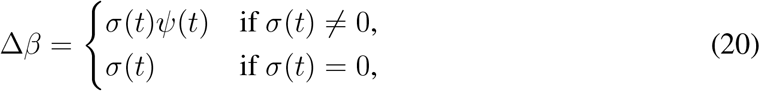

when *Ψ*(*t*) = 0, indicating no change in the motion. The change of speed of the follower is calculated by

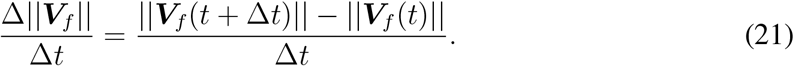

##### Relative motion between leader and follower

This section computed the relative average change in speed (Δ ∥*V*_*f*_∥) and direction of motion (Δ*β*) as a function of relative coordinates of group mates as shown in Fig. 14b. The unit vector pointing from follower *f* to leader *l* is calculated as

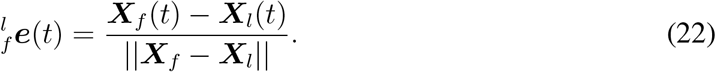

The unsigned angle between the direction of motion of follower *f* and the unit vector pointing from follower *f* to leader *l* is represented by:

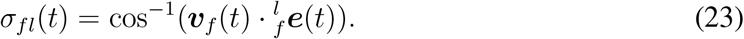

The relative direction of the motion of the follower sign can be determined as

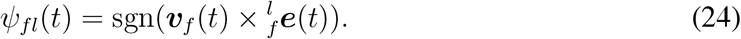

Thus, the signed change of the direction of motion is

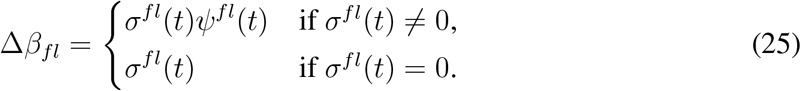

The polar coordinates are then converted to rectangular coordinates 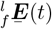 by

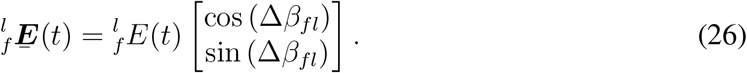

#### Force map

In this 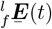 relative coordinate system, a focal individual (leader) *l* is located at the origin (0, 0), with velocity vector aligned with positive *x* axis. This local leader-centered domain was partitioned into a series of overlapping square bin regions. For each focal follower *f* and leader *l* pair, the changes of speed 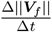 and direction of motion 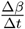 are stored in all corresponding bins. Once all the data for these measures of interest are binned, the mean value of each bin is determined. For a square domain with edge length *F*, this method divided the domain into overlapping square bins centered on each focal follower as 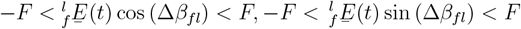 respectively (Mudaliar and Schaerf, 2020). Relative *x* and *y* coordinates describe the position within this 2D domain. The bins within the square domain are identified by paired column and row indices such that both left and right edges are separated by *q < r*. The left edge of the domain is located at − *F*, − *F* + *r*, − *F* + 2*r*, …, *F* − *q*, and the right edges is located at − *F* + *q*, − *F* + *q* + *r*, − *F* + *q* + 2*r*, …, *F*. In a similar way, the top and bottom edges can also be partitioned. The mean 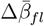 from all the leader-follower pairs’ angular differences Δ*β*_*fl*_ was calculated as

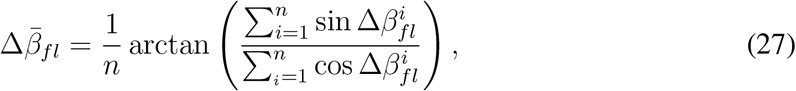

where *n* is the number of leader-follower pairs and the 4 quadrant arctangent (`>>arctan2`) function was used. Finally, the mean angle representing the alignment was visualized in a vector plot of directional arrows (`>>quiver`).

### Feedback control rule (PID)

When followers decided to re-engage the leaders, the feedback control law governing this velocity-matching behavior was investigated as illustrated in Fig. 8a. For a leader and follower with measured velocities ***V***_*l*_(*t*) and ***V***_*f*_ (*t*), a parallel simulation was constructed in which a simulated follower with velocity 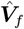 implemented velocity feedback on the proportional-integral-derivative (PID) velocity error ***E***_*r*_(*t*) between the leader and simulated follower as

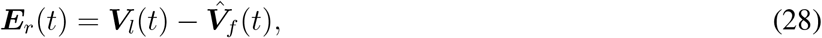

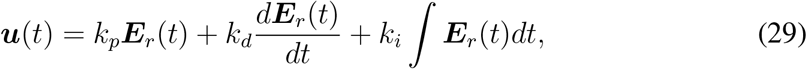

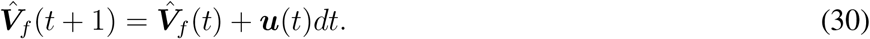

Here, ***u***(*t*) represents the control input, and *dt* is the sampling time interval. The proportional, derivative, and integral gains *k*_*p*_, *k*_*d*_, *k*_*i*_ were determined from a non-linear optimization. The optimization’s cost function is the difference between the true and simulated follower velocity and the gains *k*_*p*_, *k*_*d*_, *k*_*i*_ were found as

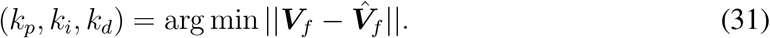

Nonlinear minimization is prone to convergence to local minima. To avoid accepting convergence to a local minimum and allow the error-minimizing gains to serve as a useful description of the behavior, gains were accepted only when the FIT percentage, calculated by

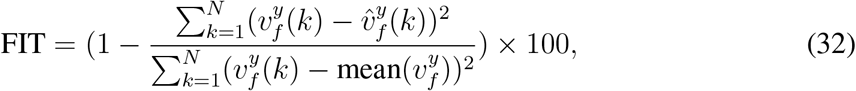

exceeded 70%.

Next, the relationship between the average inter-agent distance and the identified gains was examined. The average inter-agent distance of each trial was calculated as

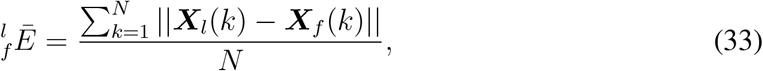

where ***X***_*l*_ and ***X***_*f*_ are the 3D positions of the leader and follower, and *N* is the total number of data points of that trial.

## Acknowledgments

This work was supported in part by IAF’s ONR Young Investigator Award N00014-19-1-2216. SI declares no potential conflict of interest.

## Author contributions statement

SI: Conceptualization, Methodology, Investigation, Software, Writing-Original draft preparation, Visualization, Writing-Reviewing and Editing. IAF: Supervision, Project administration, Funding acquisition, Conceptualization, Investigation, Writing-Original draft preparation, Visualization, Writing-Reviewing and Editing.

## Supplementary materials

### Data availability statement

The data used in this experiment is available at (link is private while this manuscript is under review).

**S1 Video. Examples of pairwise flights**. This video shows examples of honeybees pairwise motion.

**S2 Video. Examples of pairwise flights**. These videos show several examples of honeybees pairwise motion.

**S3 Video. Video of an example leader-follower behavior with tracking**. In this video the follower modified their optical expansion rate which produced three different zones.

**S4 Video. Video of an example leader-follower behavior with disengaging**. Optical expansion rate with disengaging behaviors.

**S5 Video. Video of tracking behavior**. The follower tries to match its velocity with the leader.

**S6 Video. Example leader-follower trajectories in static stimulus**. The follower insect follows the leader in static stimulus.

**S7 Video. Example leader-follower trajectories away from stimulus**. The follower insect follows the leader away from stimulus.

## Appendix

### Out-degree relative to in-degree

This study used the growth relationship of in-degree gain with group size, as seen in Fig. 3g because of its clear relationship with information flow. Some references use an alternate edge orientation, and it is helpful to note in Fig. 11 that out-degree also follows a similar growth trend with group size.

**Figure 11:**
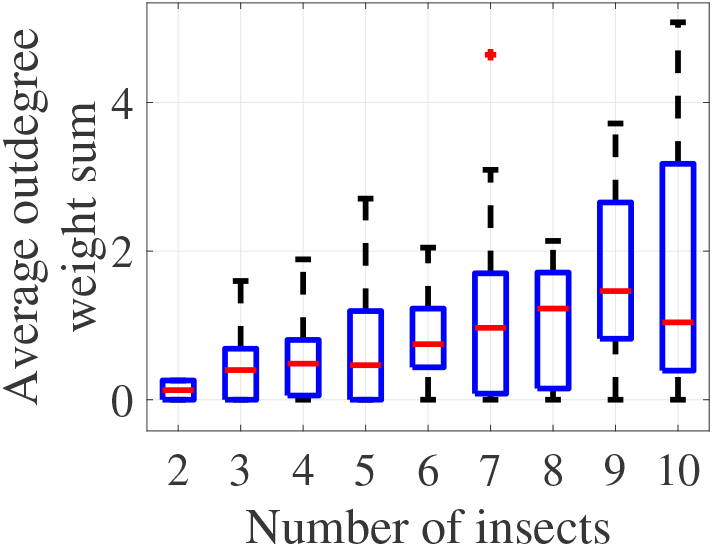
Out-degree weight sum vs. group size follows a similar trend as in-degree weight in Fig. 3g.

**Figure 12:**
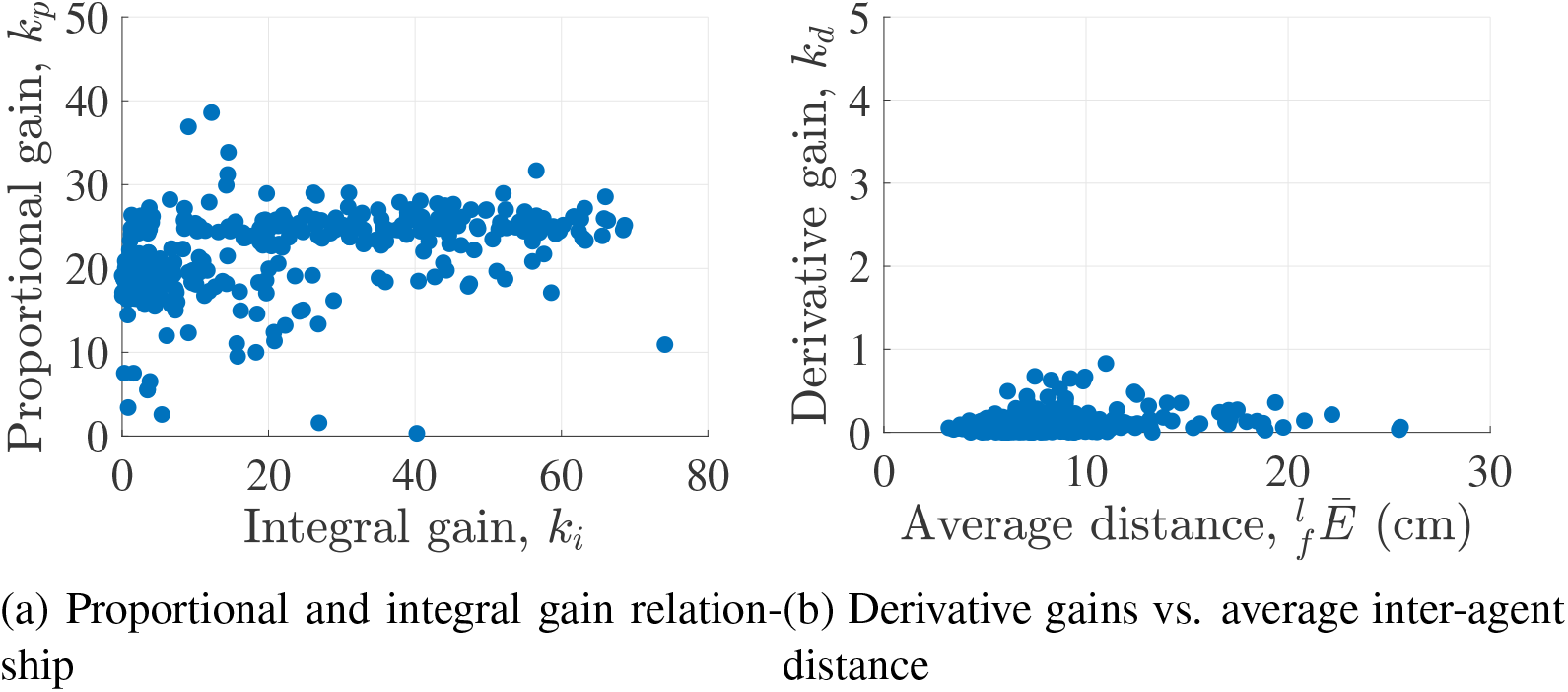
(a) Relationship between proportional and integral gains, (b) Sum of out-degree weights vs. group size, (c) Derivative gains vs. average inter-agent distance.

### Proportional, integral, and derivative gain relationships and magnitude

The identified proportional and integral gains had no clear correlation, as seen in Fig. 12a. Derivative gain was small, and negligible, as seen in Fig. 12b.

### Static stimulus case

370 trials having a static stimulus were segmented from the main dataset to identify if the overall behavior persisted without a moving stimulus. Optical expansion rate, seen in Fig. 13a, followed the same overall trend with a higher magnitude mean and variability within 10 cm inter-agent distance. Importantly, optical expansion rate was best matched by a constant beyond 10 cm, indicating that optical expansion rate regulation remains the best optical regulation candidate. Long range engagements (beyond 30 cm) were not observed in the static case. The increasing slope of velocity (Fig. 13c) with respect to inter-agent distance was preserved, while the negative slope beyond 30 cm could not be resolved due to the inflection point at 30 cm. Considering inter-agent distance in Fig. 13b, initial engagement occurs at a lower inter-agent distance in static trials, and is not as tightly regulated as time continues. Instead, a slight increase is seen in the mean trend, which does not exceed the variance.

**Figure 13:**
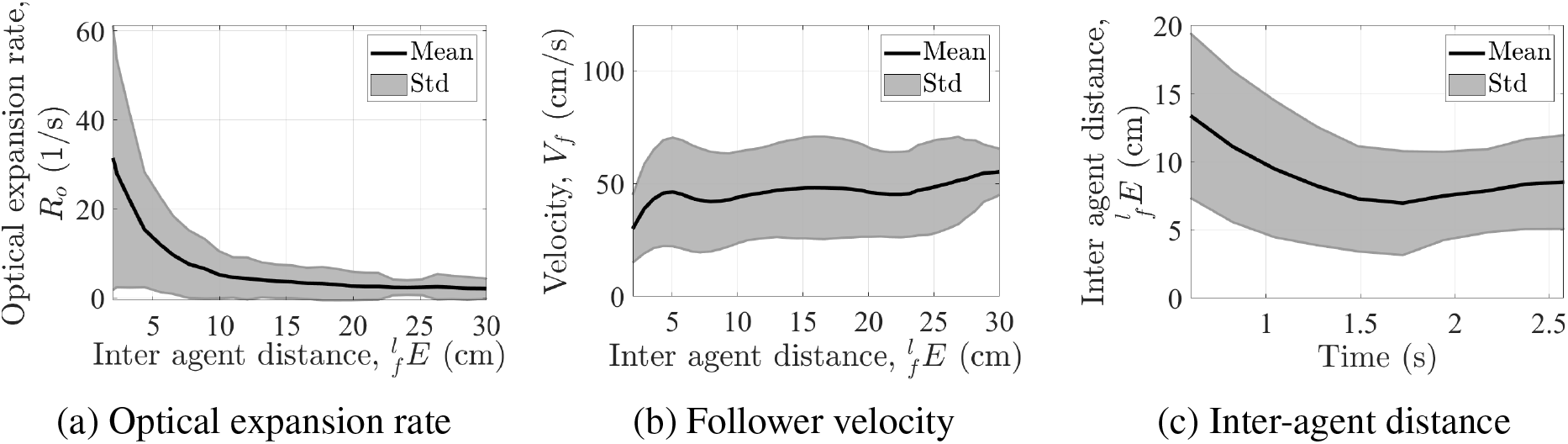
370 trials with a static stimulus were segmented to determine if the optical expansion rate (a) regulated above 10 cm, and the follower velocity (b) proportionately scaled by interagent distance whereas the inter-agent distance (c) showed an inconsistent trend.

Broadly, the static stimulus case reflects similar trends with higher variability and a lower engagement distance. In particular, the key finding of locked optical expansion rate beyond 10 cm persists.

### Heading angle analysis: detailed definitions and additional results

**Figure 14:**
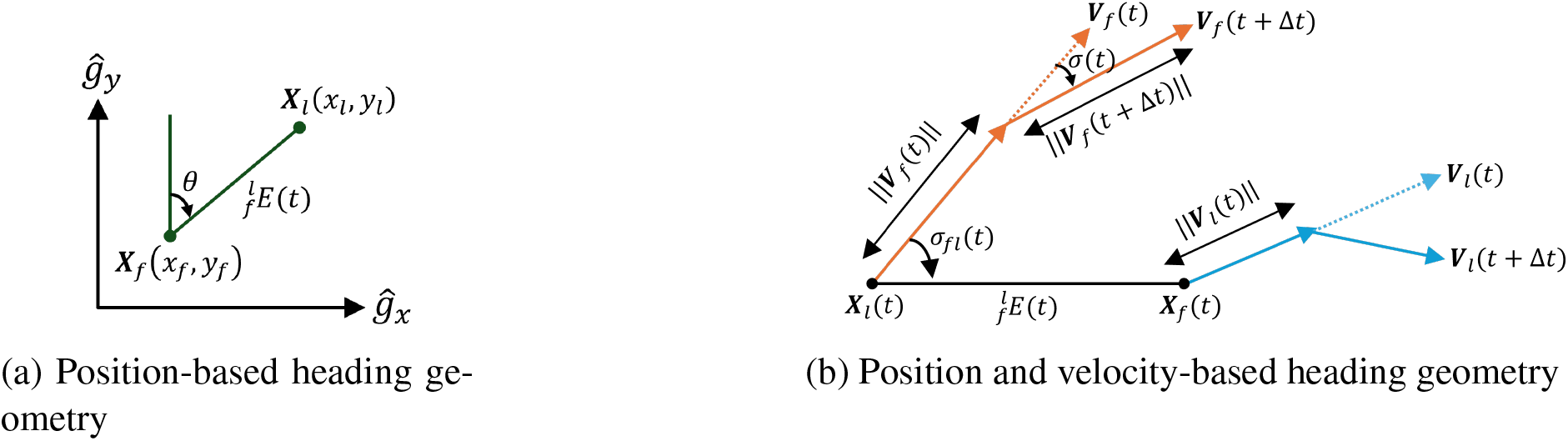
The position-based heading angle, *θ*, was calculated (a) based on the line connecting the follower and leader points ***X***_*f*_ and ***X***_*l*_. Velocity may also have effects on heading, and distributed approach operates in the follower-relative frame, thus a *relative* heading angle *σ*_*fl*_ was also computed (b) using both positions and velocities. *σ*_*fl*_ describes the angle between the follower velocity vector (orange) relative to the position vector between ***X***_*f*_ and ***X***_*l*_. The follower (orange) and leader (blue) velocity vectors are considered at both time *t* and the following timestep *t* + Δ*t*, with the change in follower velocity direction labeled *σ*(*t*).

**Figure 15:**
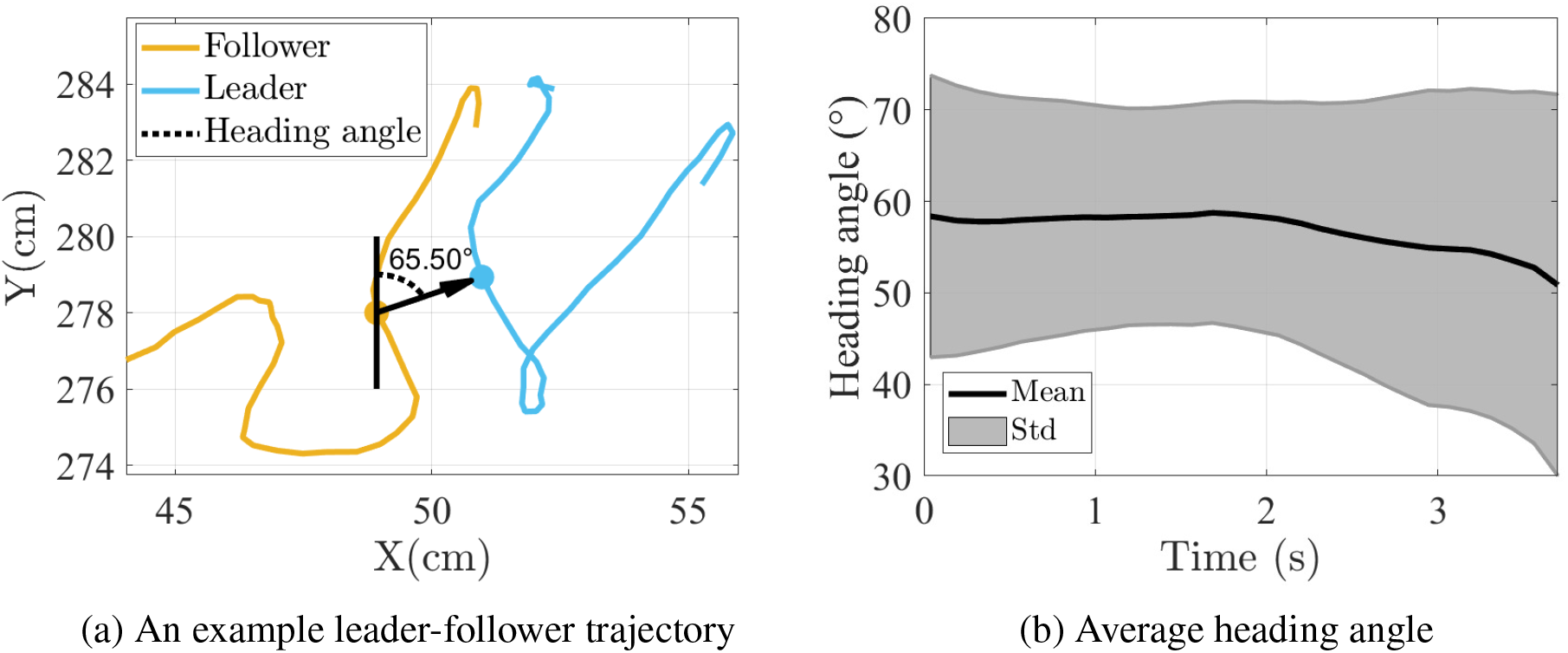
An example of leader-follower trajectories is provided to illustrate the calculated heading angle of 65.50° in (a), where each pair of points was used to determine the heading angle between the vertical line and the line joining the points, and in (b), the average heading angle across all leader-follower pairs (mean depicted in black and standard deviation in gray shading) shows a decreasing trend over time from approximately 60° to 50°.

**Figure 16:**
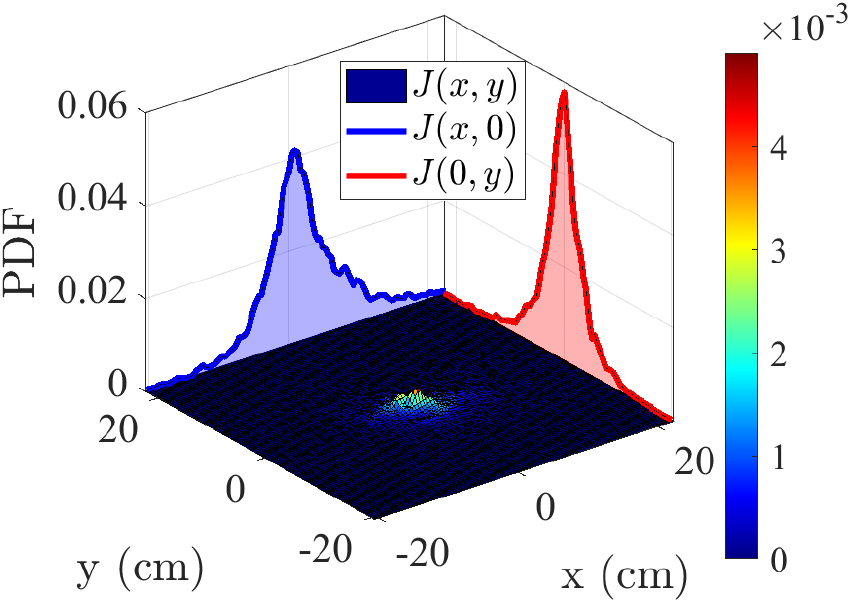
The probability density function (PDF) of the follower’s position in the relative coordinate system is represented as *J*(*x*, 0) for the x-coordinate and *J*(0, *y*) for the y-coordinate, with the joint probability density denoted as *J*(*x, y*). The PDF indicates that the followers’ relative positions were primarily clustered around the center where the leaders were positioned.

**Figure 17:**
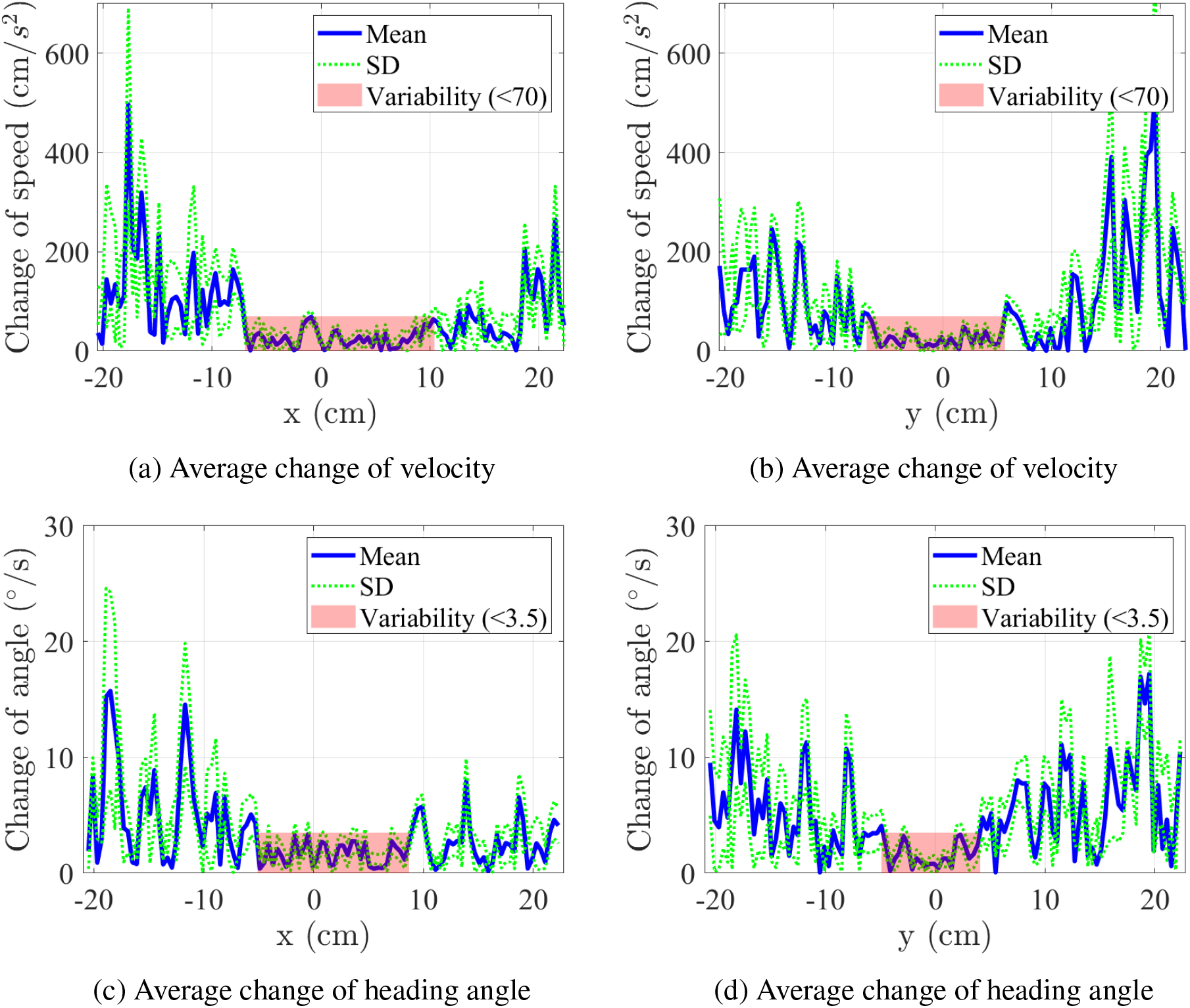
The absolute average follower’s speed change along both the *x* and *y* axes within the relative coordinate system shows lower variability when the speed is less than 70 (for *x* coordinate the distance is from −7 to 10.4 cm and for *y* coordinate the distance is from −7 to 5.7 cm), but exhibits higher variability beyond this threshold, as seen in figures (a) and (b). Similarly, the absolute change in angle shows lower variability when it is less than 3.5° with ranges of −5.1 to 8.7 cm for the *x* coordinate and −4.9 to 4.1 cm for the *y* coordinate, as shown in (c) and (d).

**Figure 18:**
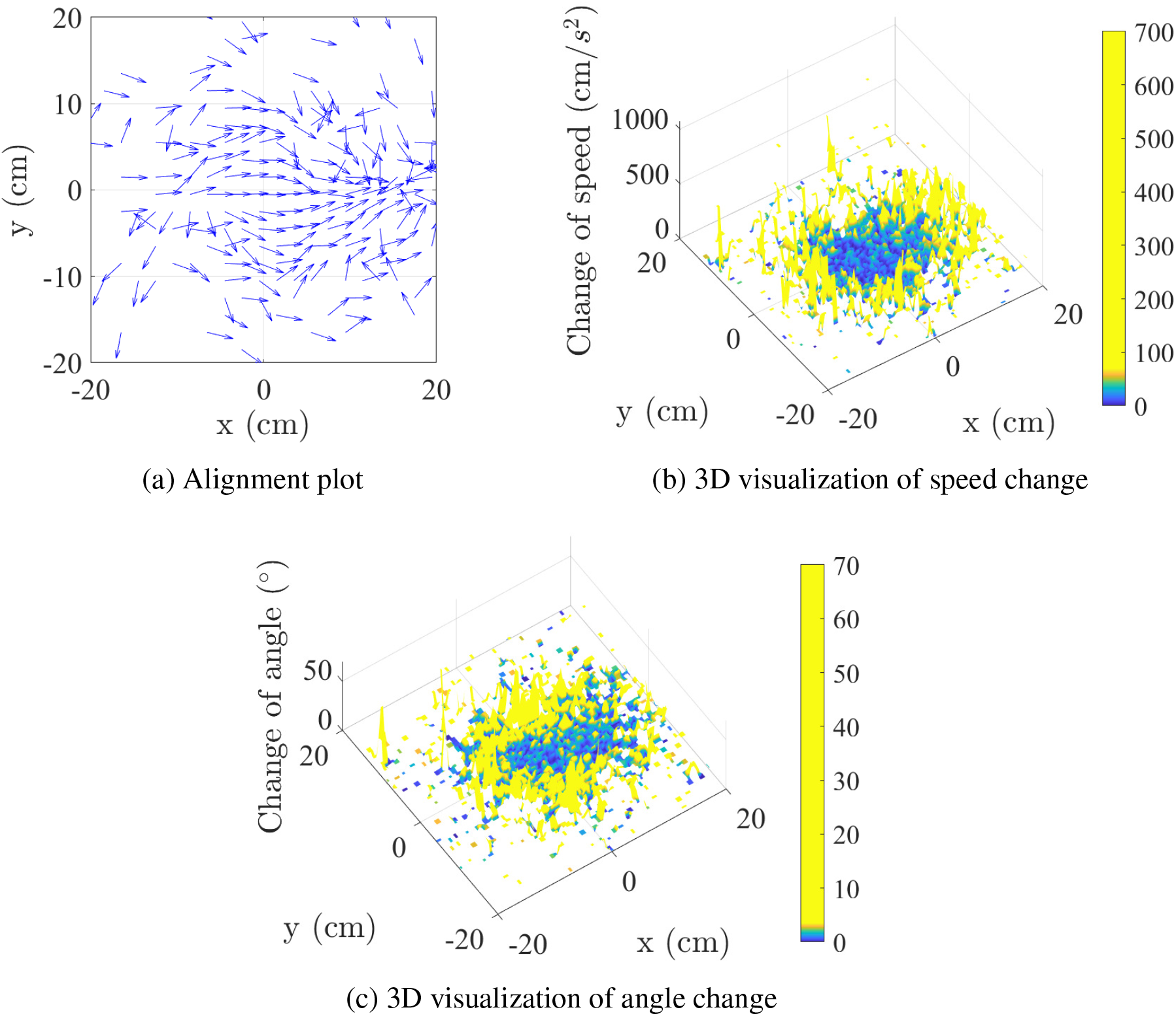
(a) The arrow depicts the average relative direction of motion of the followers in the relative coordinate system. A straighter arrow indicates a greater similarity in direction between pairs of insects. At a closer distance, the arrows become much straighter, representing a high alignment between the two insects. (b) The change in speed can also be depicted through surface plot, which is a 3D graphical representation of Fig. 17a and 17b, using a threshold of 70 to identify areas of lower variability (blue color region) at closer distances. (c) The change in angle is segmented by 3.5°, representing lower variability at closer distances. This is also shown as a 3D graphical representation of Fig. 17c and 17d.

## References

C. Agrillo, M. Dadda, G. Serena, and A. Bisazza. Do fish count? spontaneous discrimination of quantity in female mosquitofish. Animal Cognition, 11(3):495–503, Feb. 2008. doi: 10.1007/s10071-008-0140-9. URL https://doi.org/10.1007/s10071-008-0140-9.

I. Aoki. A simulation study on the schooling mechanism in fish. NIPPON SUISAN GAKKAISHI, 48(8):1081–1088, 1982. doi: 10.2331/suisan.48.1081. URL https://doi.org/10.2331/suisan.48.1081.

G. Ariel, Y. Ophir, S. Levi, E. Ben-Jacob, and A. Ayali. Individual pause-and-go motion is instrumental to the formation and maintenance of swarms of marching locust nymphs. PLoS ONE, 9(7):e101636, July 2014. doi: 10.1371/journal.pone.0101636. URL https://doi.org/10.1371/journal.pone.0101636.

A. Attanasi, A. Cavagna, L. D. Castello, I. Giardina, S. Melillo, L. Parisi, O. Pohl, B. Rossaro, E. Shen, E. Silvestri, and M. Viale. Collective behaviour without collective order in wild swarms of midges. PLoS Computational Biology, 10(7):e1003697, July 2014. doi: 10.1371/journal.pcbi.1003697. URL https://doi.org/10.1371/journal.pcbi.1003697.

H. Autrum. Electrophysiological analysis of the visual systems in insects. Exp. Cell Res., 14 (Suppl 5):426–439, 1958.

M. Ballerini, N. Cabibbo, R. Candelier, A. Cavagna, E. Cisbani, I. Giardina, V. Lecomte, A. Orlandi, G. Parisi, A. Procaccini, M. Viale, and V. Zdravkovic. Interaction ruling animal collective behavior depends on topological rather than metric distance: Evidence from a field study. Proceedings of the National Academy of Sciences, 105(4):1232–1237, 2008. doi: 10.1073/pnas.0711437105. URL https://www.pnas.org/doi/abs/10.1073/pnas.0711437105.

M. Bell. Group decision making by insects and humans. 13th International Command and Control Research and Technology, 06 2008. URL https://www.researchgate.net/publication/236345575_Group_Decision_Making_by_Insects_and_Humans.

M. A. Billah and I. A. Faruque. Visually guided swarm motion coordination via insect-inspired small target motion reactions. Bioinspiration & Biomimetics, 19(5):056013, aug 2024. doi: 10.1088/1748-3190/ad6726. URL https://dx.doi.org/10.1088/1748-3190/ad6726.

N. W. Bode, A. J. Wood, and D. W. Franks. The impact of social networks on animal collective motion. Animal Behaviour, 82(1):29–38, July 2011. doi: 10.1016/j.anbehav.2011.04.011. URL https://doi.org/10.1016/j.anbehav.2011.04.011.

N. W. F. Bode, A. J. Wood, and D. W. Franks. Social networks and models for collective motion in animals. Behavioral Ecology and Sociobiology, 65(2):117–130, Nov. 2010. doi: 10.1007/s00265-010-1111-0. URL https://doi.org/10.1007/s00265-010-1111-0.

J. Buhl, D. J. T. Sumpter, I. D. Couzin, J. J. Hale, E. Despland, E. R. Miller, and S. J. Simpson. From disorder to order in marching locusts. Science, 312(5778):1402–1406, June 2006. doi: 10.1126/science.1125142. URL https://doi.org/10.1126/science.1125142.

F. Bullo. Lectures on Network Systems. Kindle Direct Publishing, 1.7 edition, 2024. ISBN 978-1986425643. URL https://fbullo.github.io/lns.

A. Cavagna, A. Cimarelli, I. Giardina, G. Parisi, R. Santagati, F. Stefanini, and M. Viale. Scalefree correlations in starling flocks. Proceedings of the National Academy of Sciences, 107 (26):11865–11870, June 2010. doi: 10.1073/pnas.1005766107. URL https://doi.org/10.1073/pnas.1005766107.

L. Chittka. The mind of a bee. June 2022. doi: 10.1515/9780691236247. URL https://doi.org/10.1515/9780691236247.

A. K. Churchland, R. Kiani, and M. N. Shadlen. Decision-making with multiple alternatives. Nature Neuroscience, 11(6):693–702, May 2008. doi: 10.1038/nn.2123. URL https://doi.org/10.1038/nn.2123.

I. D. Couzin, J. Krause, R. James, G. D. Ruxton, and N. R. Franks. Collective memory and spatial sorting in animal groups. Journal of Theoretical Biology, 218(1):1–11, Sept. 2002. doi: 10.1006/jtbi.2002.3065. URL https://doi.org/10.1006/jtbi.2002.3065.

F. Cucker and S. Smale. Emergent behavior in flocks. IEEE Transactions on Automatic Control, 52(5):852–862, May 2007. ISSN 0018-9286. doi: 10.1109/tac.2007.895842. URL http://dx.doi.org/10.1109/TAC.2007.895842.

A. Czirók and T. Vicsek. Collective behavior of interacting self-propelled particles. Physica A: Statistical Mechanics and its Applications, 281(1):17–29, 2000. ISSN 0378-4371. doi: 10.1016/S0378-4371(00)00013-3. URL https://www.sciencedirect.com/science/article/pii/S0378437100000133.

J. A. Downes. The swarming and mating flight of diptera. Annual Review of Entomology, 14(1):271–298, 1969. doi: 10.1146/annurev.en.14.010169.001415. URL https://doi.org/10.1146/annurev.en.14.010169.001415.

B. R. Fajen and W. H. Warren. Visual guidance of intercepting a moving target on foot. Perception, 33(6):689–715, 2004. doi: 10.1068/p5236. URL https://doi.org/10.1068/p5236. PMID: 15330365.

J. Gautrais, F. Ginelli, R. Fournier, S. Blanco, M. Soria, H. Chaté, and G. Theraulaz. Deciphering interactions in moving animal groups. PLoS Computational Biology, 8(9):e1002678, Sept. 2012. doi: 10.1371/journal.pcbi.1002678. URL https://doi.org/10.1371/journal.pcbi.1002678.

K. Ghose, T. K. Horiuchi, P. S. Krishnaprasad, and C. F. Moss. Echolocating bats use a nearly time-optimal strategy to intercept prey. PLoS Biology, 4(5):e108, Apr. 2006. doi: 10.1371/journal.pbio.0040108. URL https://doi.org/10.1371/journal.pbio.0040108.

F. Ginelli and H. Chaté. Relevance of metric-free interactions in flocking phenomena. Physical Review Letters, 105(16), Oct. 2010. doi: 10.1103/physrevlett.105.168103. URL https://doi.org/10.1103/physrevlett.105.168103.

L. Giuggioli, T. J. McKetterick, and M. Holderied. Delayed response and biosonar perception explain movement coordination in trawling bats. PLOS Computational Biology, 11(3): e1004089, Mar. 2015. doi: 10.1371/journal.pcbi.1004089. URL https://doi.org/10.1371/journal.pcbi.1004089.

J. I. Gold and M. N. Shadlen. The neural basis of decision making. Annual Review of Neuroscience, 30(1):535–574, July 2007. doi: 10.1146/annurev.neuro.29.051605.113038. URL https://doi.org/10.1146/annurev.neuro.29.051605.113038.

D. Gorbonos, J. G. Puckett, K. van der Vaart, M. Sinhuber, N. T. Ouellette, and N. S. Gov. Pair formation in insect swarms driven by adaptive long-range interactions. Journal of The Royal Society Interface, 17(171):20200367, 2020. doi: 10.1098/rsif.2020.0367. URL https://royalsocietypublishing.org/doi/abs/10.1098/rsif.2020.0367.

P. Goyal, J. L. van Leeuwen, and F. T. Muijres. Bumblebees land rapidly by intermittently accelerating and decelerating toward the surface during visually guided landings. iScience, 25(5):104265, 2022. ISSN 2589-0042. doi: 10.1016/j.isci.2022.104265. URL https://www.sciencedirect.com/science/article/pii/S2589004222005351.

R. Gray, A. Franci, V. Srivastava, and N. E. Leonard. Multiagent decision-making dynamics inspired by honeybees. IEEE Transactions on Control of Network Systems, 5(2):793–806, 2018. doi: 10.1109/TCNS.2018.2796301.

M. Gries and N. Koeniger. Straight forward to the queen: pursuing honeybee drones (apis mellifera l.) adjust their body axis to the direction of the queen. Journal of Comparative Physiology A, 179:539–544, 1996.

A. Huth and C. Wissel. The simulation of the movement of fish schools. Journal of Theoretical Biology, 156(3):365–385, June 1992. doi: 10.1016/s0022-5193(05)80681-2. URL https://doi.org/10.1016/s0022-5193(05)80681-2.

M. S. Islam and I. A. Faruque. Experimental identification of individual insect visual tracking delays in free flight and their effects on visual swarm patterns. PLOS ONE, 17(11):1–23, 11 2022. doi: 10.1371/journal.pone.0278167. URL https://doi.org/10.1371/journal.pone.0278167.

L. Jiang, L. Giuggioli, A. Perna, R. Escobedo, V. Lecheval, C. Sire, Z. Han, and G. Theraulaz. Identifying influential neighbors in animal flocking. PLOS Computational Biology, 13(11):e1005822, Nov. 2017. doi: 10.1371/journal.pcbi.1005822. URL https://doi.org/10.1371/journal.pcbi.1005822.

J. W. Jolles, A. J. King, A. Manica, and A. Thornton. Heterogeneous structure in mixed-species corvid flocks in flight. Animal Behaviour, 85(4):743–750, Apr. 2013. ISSN 0003-3472. doi: 10.1016/j.anbehav.2013.01.015. URL http://dx.doi.org/10.1016/j.anbehav.2013.01.015.

D. Kahneman. Thinking, Fast and Slow. Farrar, Straus and Giroux, 2011. ISBN 9780141033570.

S. A. Khan, K. A. Khan, S. Kubik, S. Ahmad, H. A. Ghramh, A. Ahmad, M. Skalicky, Z. Naveed, S. Malik, A. Khalofah, et al. Electric field detection as floral cue in hoverfly pollination. Scientific Reports, 11(1):18781, 2021.

B. S. Lanchester and R. Mark. Pursuit and prediction in the tracking of moving food by a teleost fish (acanthaluteres spilomelanurus). Journal of Experimental Biology, 63(3):627–645, 1976.

B. Lemasson, J. Anderson, and R. Goodwin. Collective motion in animal groups from a neurobiological perspective: The adaptive benefits of dynamic sensory loads and selective attention. Journal of Theoretical Biology, 261(4):501–510, Dec. 2009. doi: 10.1016/j.jtbi.2009.08.013. URL https://doi.org/10.1016/j.jtbi.2009.08.013.

B. H. Lemasson, J. J. Anderson, and R. A. Goodwin. Motion-guided attention promotes adaptive communications during social navigation. Proceedings of the Royal Society B: Biological Sciences, 280(1754):20122003, Mar. 2013. doi: 10.1098/rspb.2012.2003. URL https://doi.org/10.1098/rspb.2012.2003.

H.-T. Lin and A. Leonardo. Heuristic rules underlying dragonfly prey selection and interception. Current Biology, 27(8):1124–1137, 2017. ISSN 0960-9822. doi: 10.1016/j.cub.2017.03.010. URL https://www.sciencedirect.com/science/article/pii/S0960982217302798.

H. Ling, G. E. Mclvor, K. van der Vaart, R. T. Vaughan, A. Thornton, and N. T. Ouellette. Costs and benefits of social relationships in the collective motion of bird flocks. Nature Ecology and Evolution, 3(6):943–948, May 2019a. doi: 10.1038/s41559-019-0891-5. URL https://doi.org/10.1038/s41559-019-0891-5.

H. Ling, G. E. Mclvor, K. van der Vaart, R. T. Vaughan, A. Thornton, and N. T. Ouellette. Costs and benefits of social relationships in the collective motion of bird flocks. Nature Ecology & Evolution, 3(6):943–948, Jun 2019b. ISSN 2397-334X. doi: 10.1038/s41559-019-0891-5. URL https://doi.org/10.1038/s41559-019-0891-5.

U. Lopez, J. Gautrais, I. D. Couzin, and G. Theraulaz. From behavioural analyses to models of collective motion in fish schools. Interface Focus, 2(6):693–707, Oct. 2012. doi: 10.1098/rsfs.2012.0033. URL https://doi.org/10.1098/rsfs.2012.0033.

W. M. Lord, J. Sun, N. T. Ouellette, and E. M. Bollt. Inference of causal information flow in collective animal behavior. IEEE Transactions on Molecular, Biological and Multi-Scale Communications, 2(1):107–116, June 2016. doi: 10.1109/tmbmc.2016.2632099. URL https://doi.org/10.1109/tmbmc.2016.2632099.

H. MaBouDi, J. A. Marshall, N. Dearden, and A. B. Barron. How honey bees make fast and accurate decisions. eLife, 12, June 2023. doi: 10.7554/elife.86176. URL https://doi.org/10.7554/elife.86176.

J. A. R. Marshall, R. Bogacz, A. Dornhaus, R. Planqué, T. Kovacs, and N. R. Franks. On optimal decision-making in brains and social insect colonies. Journal of The Royal Society Interface, 6(40):1065–1074, Feb. 2009. doi: 10.1098/rsif.2008.0511. URL https://doi.org/10.1098/rsif.2008.0511.

T. F. Mathejczyk and M. F. Wernet. Heading choices of flying drosophila under changing angles of polarized light. Scientific Reports, 9(1):16773, Nov. 2019.

A. S. Mauss and A. Borst. Optic flow-based course control in insects. Current Opinion in Neurobiology, 60:21–27, 2020.

M. K. McBeath, D. M. Shaffer, and M. K. Kaiser. How baseball outfielders determine where to run to catch fly balls. Science, 268(5210):569–573, 1995. doi: 10.1126/science.7725104. URL https://www.science.org/doi/abs/10.1126/science.7725104.

M. Mesbahi and M. Egerstedt. Graph Theoretic Methods in Multiagent Networks, volume 33. Princeton University Press, 2010.

R. K. Mudaliar and T. M. Schaerf. Examination of an averaging method for estimating repulsion and attraction interactions in moving groups. PLOS ONE, 15(12):1–28, 12 2020. doi: 10.1371/journal.pone.0243631. URL https://doi.org/10.1371/journal.pone.0243631.

P. R. Murphy, E. Boonstra, and S. Nieuwenhuis. Global gain modulation generates time-dependent urgency during perceptual choice in humans. Nature Communications, 7 (1), Nov. 2016. doi: 10.1038/ncomms13526. URL https://doi.org/10.1038/ncomms13526.

G. Nagy, A. Thornton, H. Ling, G. McIvor, N. T. Ouellette, and R. Vaughan. Computational and structural advantages of pairwise flocking. In 2019 International Symposium on Multi-Robot and Multi-Agent Systems (MRS). IEEE, Aug. 2019. doi: 10.1109/mrs.2019.8901049. URL https://doi.org/10.1109/mrs.2019.8901049.

M. Nagy, G. Vásárhelyi, B. Pettit, I. Roberts-Mariani, T. Vicsek, and D. Biro. Context-dependent hierarchies in pigeons. Proceedings of the National Academy of Sciences, 110 (32):13049–13054, July 2013. doi: 10.1073/pnas.1305552110. URL https://doi.org/10.1073/pnas.1305552110.

R. M. Neems, J. Lazarus, and A. J. Mclachlan. Swarming behavior in male chironomid midges: a cost-benefit analysis. Behavioral Ecology, 3(4):285–290, 1992. doi: 10.1093/beheco/3.4.285. URL https://doi.org/10.1093/beheco/3.4.285.

A. Okubo and H. C. Chiang. An analysis of the kinematics of swarming of anarete pritchardi kim (diptera: Cecidomyiidae). Population Ecology, 16(1):1–42, Sept. 1974. doi: 10.1007/bf02514077. URL https://doi.org/10.1007/bf02514077.

R. M. Olberg, A. H. Worthington, and K. R. Venator. Prey pursuit and interception in dragonflies. Journal of Comparative Physiology A, 186(2):155–162, Feb 2000. ISSN 1432-1351. doi: 10.1007/s003590050015. URL https://doi.org/10.1007/s003590050015.

A. Papana. Connectivity analysis for multivariate time series: Correlation vs. causality. Entropy, 23(12):1570, Nov. 2021. doi: 10.3390/e23121570. URL https://doi.org/10.3390/e23121570.

A. Paz, K. J. Holt, A. Clarke, A. Aviles, B. Abraham, A. C. Keene, E.R. Duboué, Y. Fily, and J. E. Kowalko. Changes in local interaction rules during ontogeny underlie the evolution of collective behavior. iScience, 26(9):107431, 2023. ISSN 2589-0042. doi: 10.1016/j.isci.2023.107431. URL https://www.sciencedirect.com/science/article/pii/S2589004223015080.

J. G. Puckett, R. Ni, and N. T. Ouellette. Time-frequency analysis reveals pairwise interactions in insect swarms. Physical Review Letters, 114(25), June 2015. doi: 10.1103/physrevlett.114.258103. URL https://doi.org/10.1103/physrevlett.114.258103.

M. B. Reiser and M. H. Dickinson. Drosophila fly straight by fixating objects in the face of expanding optic flow. Journal of Experimental Biology, 213(10):1771–1781, 2010.

C. W. Reynolds. Flocks, herds and schools: A distributed behavioral model. New York, NY, USA, 1987. Association for Computing Machinery. ISBN 0897912276. doi: 10.1145/37401.37406. URL https://doi.org/10.1145/37401.37406.

L. Ristroph, G. Ristroph, S. Morozova, A. J. Bergou, S. Chang, J. Guckenheimer, Z. J. Wang, and I. Cohen. Active and passive stabilization of body pitch in insect flight. J R Soc Interface, 10(85):20130237, May 2013.

T. Schreiber. Measuring information transfer. Physical Review Letters, 85(2):461–464, July 2000. doi: 10.1103/physrevlett.85.461. URL https://doi.org/10.1103/physrevlett.85.461.

J. R. Serres and F. Ruffier. Optic flow-based collision-free strategies: From insects to robots. Arthropod structure & development, 46(5):703–717, 2017.

C. E. Shannon. A mathematical theory of communication. The Bell System Technical Journal, 27:379–423, 1948. URL http://plan9.bell-labs.com/cm/ms/what/shannonday/shannon1948.pdf.

M. V. Srinivasan and S.-W. Zhang. Visual navigation in flying insects. International review of neurobiology, 44:67–92, 2000.

J. Sun and E. M. Bollt. Causation entropy identifies indirect influences, dominance of neighbors and anticipatory couplings. Physica D: Nonlinear Phenomena, 267:49–57, Jan. 2014. doi: 10.1016/j.physd.2013.07.001. URL https://doi.org/10.1016/j.physd.2013.07.001.

J. Sun, D. Taylor, and E. M. Bollt. Causal network inference by optimal causation entropy. 2014. doi: 10.48550/ARXIV.1401.7574. URL https://arxiv.org/abs/1401.7574.

J. Talley, S. Pusdekar, A. Feltenberger, N. Ketner, J. Evers, M. Liu, A. Gosh, S. E. Palmer, T. J. Wardill, and P. T. Gonzalez-Bellido. Predictive saccades and decision making in the beetle-predating saffron robber fly. Current Biology, June 2023. doi: 10.1016/j.cub.2023.06.019. URL https://doi.org/10.1016/j.cub.2023.06.019.

K. Tan, Z. Wang, H. Li, S. Yang, Z. Hu, G. Kastberger, and B. P. Oldroyd. An ‘i see you’prey– predator signal between the asian honeybee, apis cerana, and the hornet, vespa velutina. Animal Behaviour, 83(4):879–882, 2012.

K. Tan, Z. Wang, W. Chen, Z. Hu, and B. P. Oldroyd. The ‘i see you’prey–predator signal of apis cerana is innate. Naturwissenschaften, 100:245–248, 2013.

S. E. Tavares. A comparison of integration and low-pass filtering. IEEE Transactions on Instrumentation and Measurement, 15(1/2):33–38, 1966. doi: 10.1109/TIM.1966.4313498.

D. Thura, J. Beauregard-Racine, C.-W. Fradet, and P. Cisek. Decision making by urgency gating: theory and experimental support. Journal of Neurophysiology, 108(11):2912–2930, Dec. 2012. doi: 10.1152/jn.01071.2011. URL https://doi.org/10.1152/jn.01071.2011.

M. Ulyshen, C. R. Traylor, and B. N. Danforth. Patterns of nest-site selection by colletes thoracicus within a forested watershed. Apidologie, 54(6):56, Nov 2023. ISSN 1297-9678. doi: 10.1007/s13592-023-01035-7. URL https://doi.org/10.1007/s13592-023-01035-7.

J. T. Vance, I. Faruque, and J. S. Humbert. Kinematic strategies for mitigating gust perturbations in insects. Bioinspir Biomim, 8(1):016004, Jan. 2013.

L. Varennes, H. G. Krapp, and S. Viollet. Two pursuit strategies for a single sensorimotor control task in blowfly. Scientific Reports, 10(1):1–13, 2020a. ISSN 20452322. doi: 10.1038/s41598-020-77607-9. URL https://doi.org/10.1038/s41598-020-77607-9.

L. Varennes, H. G. Krapp, and S. Viollet. Two pursuit strategies for a single sensorimotor control task in blowfly. Scientific Reports, 10(1):20762, Nov. 2020b.

T. Vicsek and A. Zafeiris. Collective motion. Physics Reports, 517(3-4):71–140, Aug. 2012. doi: 10.1016/j.physrep.2012.03.004. URL https://doi.org/10.1016/j.physrep.2012.03.004.

T. J. Wardill, S. T. Fabian, A. C. Pettigrew, D. G. Stavenga, K. Nordström, and P. T. Gonzalez-Bellido. A novel interception strategy in a miniature robber fly with extreme visual acuity. Current Biology, 27(6):854–859, Mar. 2017. doi: 10.1016/j.cub.2017.01.050. URL https://doi.org/10.1016/j.cub.2017.01.050.

T. L. Warren, P. T. Weir, and M. H. Dickinson. Flying Drosophila melanogaster maintain arbitrary but stable headings relative to the angle of polarized light. Journal of Experimental Biology, 221(9):jeb177550, 05 2018. ISSN 0022-0949. doi: 10.1242/jeb.177550. URL https://doi.org/10.1242/jeb.177550.

S. C. Whitehead, S. Leone, T. Lindsay, M. R. Meiselman, N. J. Cowan, M. H. Dickinson, N. Yapici, D. L. Stern, T. Shirangi, and I. Cohen. Neuromuscular embodiment of feedback control elements in drosophila flight. Science Advances, 8(50), Dec. 2022. doi: 10.1126/sciadv.abo7461. URL https://doi.org/10.1126/sciadv.abo7461.

M. Zumaya, H. Larralde, and M. Aldana. Delay in the dispersal of flocks moving in unbounded space using long-range interactions. Scientific Reports, 8, 10 2018. doi: 10.1038/s41598-018-34208-x.

